# Efficient variance components analysis across millions of genomes

**DOI:** 10.1101/522003

**Authors:** Ali Pazokitoroudi, Yue Wu, Kathryn S. Burch, Kangcheng Hou, Aaron Zhou, Bogdan Pasaniuc, Sriram Sankararaman

## Abstract

Variance components analysis has emerged as a powerful tool in complex trait genetics, with applications ranging from heritability estimation to association mapping. While the application of these methods to large-scale genetic datasets can potentially reveal important insights into genetic architecture, existing methods for fitting variance components do not scale well to these datasets. Here, we present a new algorithm for variance components analysis that is accurate and highly efficient, capable of estimating one hundred variance components on a million individuals genotyped at a million SNPs in a few hours. We illustrate the utility of our method in estimating variation in a trait explained by genotyped SNPs (SNP heritability) as well in partitioning heritability across population and functional genomic annotations. Analyzing 22 diverse traits with genotypes from 300, 000 individuals across about 8 million common and low frequency SNPs (minor allele frequency > 0.1%), we observe that the allelic effect size increases with decreasing MAF (minor allele frequency) and LD (linkage disequilibrium) across the analyzed traits consistent with the action of negative selection. Partitioning heritability across 28 functional annotations, we observe enrichment of heritability in FANTOM5 enhancers in asthma, eczema, thyroid and autoimmune disorders.

## Introduction

Variance components analysis [1] has emerged as a versatile tool in human complex trait genetics, enabling studies of the genetic contribution to variation in a trait [2] as well as its distribution across genomic loci [3, 4], allele frequencies [3], and functional annotations [3, 5, 6]. There is increasing interest in applying methods for variance components analysis to large-scale genetic datasets with the goal of uncovering novel insights into the genetic architecture of complex traits[7, 4]. A prominent example of the utility of these methods is in the estimation of SNP heritability 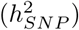 [2], the variance in a trait explained by a given set of genotyped SNPs. Variance components methods for estimating SNP heritability typically assume a genetic variance component that represents the fraction of phenotypic variation explained by the SNPs included in the study and a residual variance component. Recent studies have shown that these “single-component” methods yield biased estimates of SNP heritability due to the LD and MAF dependent architecture of complex traits [8, 9]. On the other hand, flexible models with multiple variance components [3, 4] that allows for SNP effects to vary with MAF and LD, have been shown to yield more accurate SNP heritability estimates [8, 9]. Recent work has shown that SNP heritability can be estimated with minimal assumptions about the genetic architecture [10]; however, this method cannot partition heritablity across categories of SNPs of interest such as functional or population genomic annotations. Partitioning heritability requires fitting multiple variance components, thus creating the need for accurate and scalable methods that can fit tens or even hundreds of variance components to large-scale genomic data to obtain accurate and novel insights into genetic architecture.

While the ability to fit flexible variance component models to large-scale datasets is essential to obtain accurate and novel insights into genetic architecture, fitting such models requires scalable algorithms. Approaches for estimating variance components typically search for parameter values that maximize the likelihood or the restricted maximum likelihood (REML) [11]. Despite a number of algorithmic improvements [2, 4, 12, 13, 14, 15, 16], computing REML estimates of the variance components on data sets such as the UK Biobank [17] (≈ 500, 000 individuals genotyped at nearly one million SNPs) remains challenging. The reason is that methods for computing these estimators typically perform repeated computations on the input genotypes.

We propose a new method that can jointly estimate multiple variance components efficiently. Our proposed method, RHE-mc, is a randomized multi-component version of the classical Haseman-Elston regression for heritability estimation [18, 19]. RHE-mc builds on our previously proposed method, RHE-reg [20], which uses a randomized algorithm to estimate a single variance component. RHE-mc can simultaneously estimate multiple variance components as well as estimate variance components associated with continuous annotations and overlapping annotations. Unlike RHE-reg, RHE-mc uses a nonparametric block jackknife to estimate standard errors with little computational overhead. Further, unlike REML estimation algorithms, RHE-mc requires only a single pass over the input genotypes that results in a highly memory efficient implementation. The resulting computational efficiency permits RHE-mc to jointly fit 300 variance components in less than an hour on a dataset of about 300, 000 individuals and 500, 000 SNPs, about two orders of magnitude faster than state-of-the-art methods. On a dataset of one million individuals and one million SNPs, RHE-mc can fit 100 variance components in about 12 hours.

To demonstrate its utility, we first show that RHE-mc can accurately estimate genome-wide and partitioned SNP heritability under realistic genetic architectures (the functional dependence of SNP effect sizes on MAF and LD). We applied RHE-mc to 22 traits measured across 291, 273 individuals genotyped at 459, 792 common SNPs (MAF> 1%) in the UK Biobank to obtain estimates of genome-wide SNP heritability. We then used RHE-mc to partition heritability for the 22 traits across seven million imputed SNPs (MAF > 0.1%) into 144 bins defined based on MAF and LD. We observe that the allelic effect size tends to increase with lower MAF and LD across the traits considered. Finally, we partitioned heritability for SNPs with MAF > 0.1% across 28 functional annotations. We recover previously reported enrichment of heritability in annotations corresponding to conserved regions [7] and also document enrichment of heritability in FANTOM5 enhancers in eczema, asthma, autoimmune disorders, and thyroid disorders.

## Results

### Methods overview

RHE-mc aims to fit a variance components model that relates phenotypes ***y*** measured across *N* individuals to their genotypes over *M* SNPs ***X***:

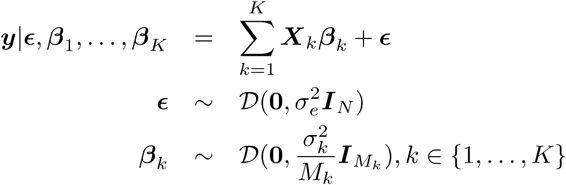

where *𝒟* (***µ, σ***^2^) is an arbitrary distribution with mean ***µ*** and variance ***σ***^2^. Each of the *M* SNPs is assigned to one of *K* non-overlapping categories so that ***X***_*k*_ is the *N × M*_*k*_ matrix consisting of standardized genotypes of SNPs belonging to category *k* (note that the expected heritability is constant within categories when we use standardized genotypes). ***β***_*k*_ denotes the effect sizes of SNPs assigned to category *k* which are drawn from a zero-mean normal distribution with variance parameter 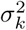 (the variance component of category *k*) while 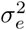 is the residual variance.

In this model, the genome-wide SNP heritability is defined as: 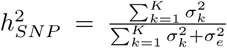 while the SNP heritability of category *k* is defined as: 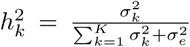. By choosing categories to represent genomic annotations of interest, *e*.*g*., chromosomes, allele frequencies, and functional annotations, these models can be used to estimate the phenotypic variation that can be attributed to the relevant annotation.

The key inference problem in this model is the estimation of the variance components: 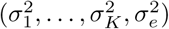. These parameters are typically estimated by maximizing the likelihood or the restricted likelihood. Instead, RHE-mc uses a scalable method-of-moments estimator, *i*.*e*., finding values of the variance components such that the population moments match the sample moments [18, 21, 22, 23, 19]. RHE-mc uses a randomized algorithm that avoids explicitly computing *N* × *N* genetic relatedness matrices that are required by method-of-moments estimators. Instead, it operates on a smaller matrix formed by multiplying the input genotype matrix with a small number of random vectors (Methods). The application of a randomized algorithm for SNP heritability estimation using a single variance component was proposed in our previous work, RHE-reg [20]. RHE-mc extends our previous work in several directions. RHE-mc can efficiently fit multiple variance components (both non-overlapping and overlapping) and can also handle continuous annotations. The resulting algorithm has scalable runtime as it only requires operating on the genotype matrix one time. Further, RHE-mc uses a streaming implementation that does not require all the genotypes to be stored in memory leading to scalable memory requirements (Supplementary Information S3). Finally, RHE-mc uses an efficient implementation of a block Jackknife to estimate standard errors with little computational overhead (Supplementary Information S1).

### Accuracy of genome-wide SNP heritability estimates in simulations

We assessed the accuracy of RHE-mc in estimating genome-wide SNP heritability as previous attempts at estimating SNP heritability have been shown to be sensitive to assumptions about how SNP effect size varies with MAF and LD[8]. Starting with genotypes of *M* = 593, 300 array SNPs over *N* = 337, 205 unrelated white British individuals in the UK Biobank, we simulated phenotypes according to 64 MAF and LD-dependent architectures by varying the SNP heritability, the proportion of variants that have non-zero effects (causal variants or CVs), the distribution of causal variants across minor allele frequencies (CVs distributed across all minor allele frequency bins or CVs restricted to either common or low-frequency bins), and the form of coupling between the SNP effect size and MAF as well as LD. For RHE-mc, we partitioned the SNPs into 24 variance components based on 6 MAF bins as well as 4 LD bins (see Methods). The key parameter in applying RHE-mc is the number of random vectors *B* which we set to 10. RHE-mc estimates were relatively insensitive when we increased the number of random vectors *B* to 100 (Supplementary Figure S1, Figure S2, Table S2)). Across these 64 architectures, RHE-mc is relatively unbiased (a test of the hypothesis of no bias is not rejected across any of the architectures at a p-value < 0.05) with the largest relative bias observed to be 0.5% of the true SNP heritability (Supplementary Figure S5). We used a block Jackknife (number of blocks = 100) to estimate the standard errors of RHE-mc and confirmed that the estimated standard errors are close to the true SE (Supplementary Table S1).

We compared the accuracy of RHE-mc to state-of-the-art methods for heritability estimation that can be applied to large datasets (across architectures where the true SNP heritability was fixed at 0.5). These methods, LDSC [24], SumHer [25], S-LDSC [26], and GRE [10], all leverage summary statistics while RHE-mc requires individual genotype data. We found that estimates from the summary-statistic methods tend to be sensitive to the underlying genetic architecture: across 16 architecture relative biases range from − 31% to 27% for LDSC, − 27% to 5% for S-LDSC, and − 5% to 9% for SumHer (Figure 1). We also compared to a recently proposed method (GRE [10]) that only estimates genome-wide SNP heritability (without partitioning by MAF/LD) and observed that our approach is as accurate as the GRE in its genome-wide estimates (relative bias of GRE ranged from 1% to 1.4%). We also considered architectures in which only rare variants are causal and found RHE-mc is accurate relative to other methods (Supplementary Figure S6). These results further emphasize that RHE-mc can accurately estimate SNP-heritability through fitting multiple variance components.

**Figure 1:**
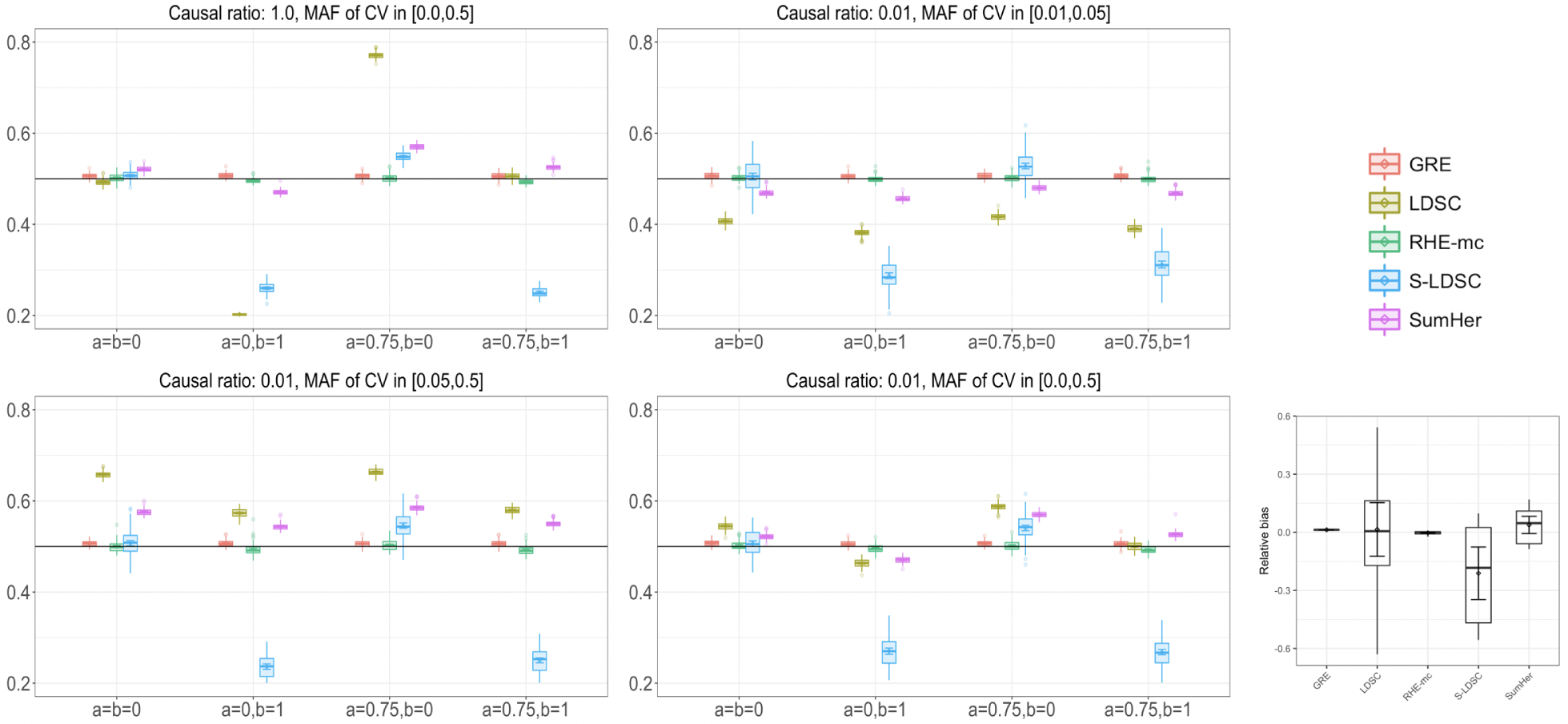
Comparison of estimates of genome-wide SNP heritability from RHE-mc with LDSC, GRE, S-LDSC, and SumHer in large-scale simulations (*N* = 337, 205 unrelated individuals, *M* = 593, 300 array SNPs). We compared methods for heritability estimation under 16 different genetic architectures. We set true heritability to 0.5 and varied the MAF range of causal variants (MAF of CV), the coupling of MAF with effect size (*a* = 0 indicates no coupling of MAF and *a* = 0.75 indicates coupling of MAF), and the effect of local LD on effect size (*b* = 0 indicates no LDAK weights and *b* = 1 indicates LDAK weights) (see Methods). Right: Relative bias of each method (as a percentage of the true *h*^2^) across 16 distinct MAF- and LD-dependent architectures. Each boxplot contains 16 points; each point represents the average estimated *h*^2^ from 100 simulations under a single genetic architecture. Here, we run RHE-mc using 24 bins formed by the combination of 6 bins based on MAF as well as 4 bins based on quartiles of the LDAK score of a SNP (see Methods). We run S-LDSC with 10 MAF bins (see Supplementary Table S5). To do a fair comparison, for every method, we computed LD scores and LDAK weights by using in-sample LD, and in all simulations we aim to estimate the SNP-heritability explained by the same set of M SNPs.

We compared RHE-mc to the state-of-the-art REML-based variance component estimation method, GCTA-mc (multi-component GREML [27, 8, 28]) and to exact multi-component Haseman-Elston Regression (HE-mc) as implemented in GCTA[27]. We ran each of these methods by partitioning SNPs into 24 variance components (6 MAF bins by 4 LD bins, see Methods). To make these experiments computationally feasible, we simulated phenotypes starting from a smaller set of genotypes (*M* = 593, 300 array SNPs and *N* = 10, 000 white British individuals). Across 16 architectures where the true SNP heritability was fixed at 0.25, the relative biases for RHE-mc range from − 3.2% to 3.6%, and from − 3.2% to 5% for GCTA-mc (Figure 2). On average, RHE-mc has standard errors that are 1.1 times larger than GCTA-mc (which range from 0.97 to 1.24) and 1.08 times larger than HE-mc (which range from 1.00 to 1.21).

**Figure 2:**
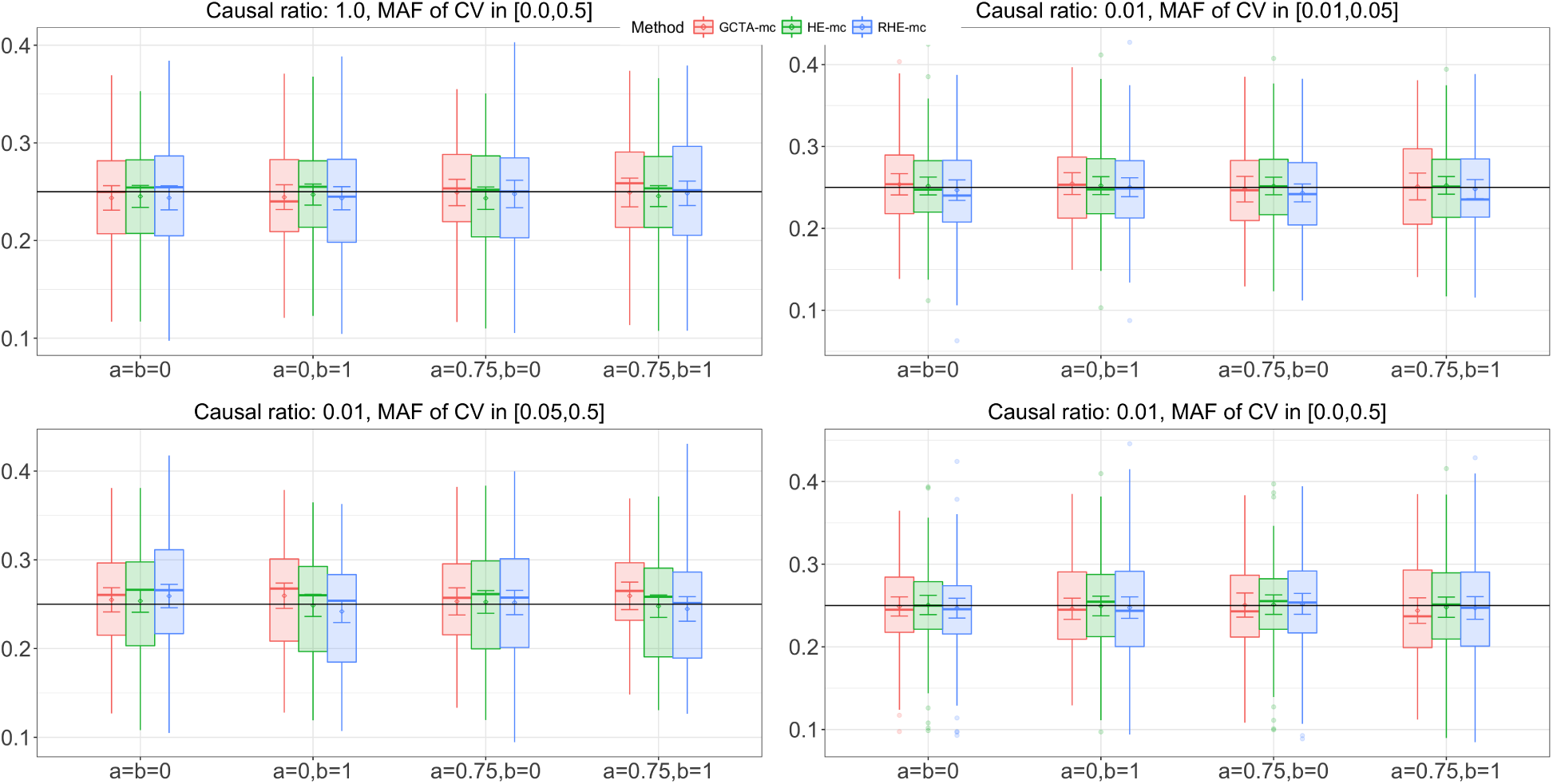
Comparison of SNP heritability estimates from RHE-mc with GCTA-mc (GCTA with multiple variance components) and HE-mc(HE with multiple variance components) (*N* = 10, 000 unrelated individuals, *M* = 593, 300 array SNPs). We compared heritability estimates from these methods under 16 different genetic architectures. We varied the MAF range of causal variants (MAF of CV), the coupling of MAF with effect size (*a*), and the effect of local LD on effect size (*b* = 0 indicates no LDAK weights and *b* = 1 indicates LDAK weights (see Methods). We ran 100 replicates where the true heritability of the phenotype is 0.25. We run RHE-mc, HE-mc and GCTA-mc using 24 bins formed by the combination of 6 bins based on MAF as well as 4 bins based on quartiles of the LDAK score of a SNP (see Methods). Across all different genetic architectures, the relative biases range from − 3.2% to 3.6% for RHE-mc, and from − 3.2% to 5% for GCTA-mc, and from − 2.6% to 1.45% for HE-mc. On average, RHE-mc has SEs that are 1.1 and 1.08 times larger than GCTA-mc and HE-mc respectively. The SE’s are computed from 100 simulations (Note that GCTA-mc did not run successfully on all 100 simulations).

### Accuracy of heritability partitioning in simulations

We also evaluated the accuracy of RHE-mc in partitioning SNP heritability in both small-scale (*M* = 593, 300 SNPs, *N* = 10, 000 individuals) (Supplementary Figure S3) and large-scale settings (*M* = 593, 300 SNPs, *N* = 337, 205 individuals) (see Supplementary Figure S4). For these experiments, we restrict our attention to architectures for which the causal variants (CVs) are chosen to lie within a narrow range of MAF. Since the variance components correspond to bins of MAF and LD, a subset of the variance components would have no causal SNPs and hence have a heritability of zero. We assess the accuracy of estimates of heritability aggregated over these components (termed the non-causal bin) as well as the heritability aggregated over the remaining genetic components (termed the causal bin). For example, variance components that correspond to MAF ∈ [0.01, 0.05] would be included in the causal bin for an architecture that restricts the MAF of CVs to lie in the range [0.01, 0.05]. For the small-scale simulations, we compared RHE-mc to GCTA-mc. We ran both methods by partitioning the SNPs into 24 variance components based on 6 MAF bins as well as 4 LD bins defined by quartiles of the measure of LDAK weight at a SNP (see Methods). Across the genetic architectures tested, estimates of heritability within each of the causal and non-causal bins are highly concordant between RHE-mc and GCTA-mc (Supplementary Figure S3, Supplementary Table S3): for the causal bin, the relative bias ranges from − 4% to 0.4% for RHE-mc and − 3.6% to 2% for GCTA-mc while, for the non-causal bin, the bias ranges from 0 to 0.7% for RHE-mc and 0 to 1.4% for GCTA-mc (Supplementary Table S3). For the large-scale settings, RHE-mc remains accurate: the relative bias ranges from − 2.6% to 3.2% (causal bin) and the bias ranges from − 0.5% to 0.2% (non-causal bin) over the genetic architectures considered (Supplementary Figure S4, Supplementary Table S4).

Heritability partitioning has been used to estimate heritability attributed to functional genomic annotations [7]. However, some of these annotations (such as FANTOM5 enhancers) are quite small covering < 1% of the genome. We explored the ability of RHE-mc to accurately estimate heritability as a function of the size of the annotation. To this end, we performed simulations using *N* = 291, 273 unrelated white British individuals and *M* = 459, 792 common SNPs. We defined 8 annotations (4 MAF bins and 2 LD bins) in which we fixed the enrichment of a selected bin and varied the proportion of SNPs in the selected category. RHE-mc obtained accurate estimates of enrichment even when the selected bin only contained 0.4% of the genome-wide SNPs (comparable to the size of FANTOM5 enhancers). RHE-mc estimates are well-calibrated: when the bin has zero enrichment, RHE-mc rejected the null hypothesis of no enrichment in 5% of the simulations while attaining high power to reject the null hypothesis even when the bin contained < 1% of the SNPs (Supplementary Information S9).

### Computational Efficiency

We benchmarked the runtime and memory usage of RHE-mc as a function of number of individuals, SNPs and variance components (Figure 3). We ran RHE-mc with *B* = 10 random vectors and 22 variance components where each chromosome forms a distinct component. On a dataset of ≈ 300, 000 individuals and ≈ 500, 000 SNPs, RHE-mc can fit 22 variance components in less than an hour and ≈ 300 variance components (corresponding to bins of size 10 Mb) with little increase in its runtime. On a dataset of one million individuals and one million SNPs, RHE-mc can fit 100 variance components in a few hours. Further, due to its use of a streaming implementation that only requires the genotypes to be operated on once, the memory requirement of RHE-mc is modest: all experiments required less than 60 GB. We compared the run time and memory usage of RHE-mc with REML-based methods (GCTA [27] and BOLT-REML [4]) on the UK Biobank genotypes consisting of around 500, 000 SNPs over varying sample sizes and observed that RHE-mc achieves several orders-of-magnitude reduction in runtime. Summary-statistic methods such as S-LDSC requires pre-computed inputs which depend on the runtimes of other softwares making a direct comparison of speed difficult. Thus, we have restricted our comparison to individual-level methods where the benchmarking can be done in a comparable manner

**Figure 3:**
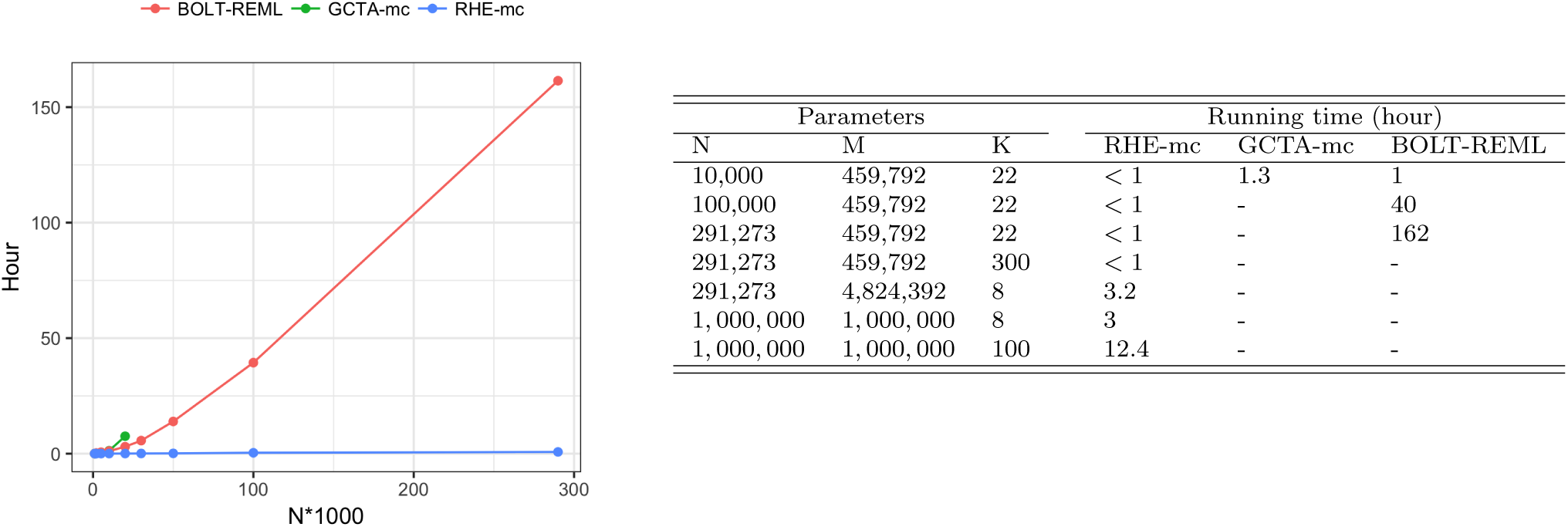
Comparison of running time of RHE-mc, GCTA-mc, and BOLT-REML. *M, N* and *K* are the number of SNPs, individuals and variance components respectively. We compared runtime of RHE-mc, GCTA-mc, and BOLT-REML with increasing sample size *N* (for a fixed number of SNPs *M* = 459, 792 and components *K* = 22). Right: RHE-mc can run efficiently even on datasets with one million individuals and SNPs as well as efficiently computing hundreds of variance components. All comparisons were performed on an Intel(R) Xeon(R) CPU 2.10 GHz server with 128 GB RAM.

### Estimating total SNP heritability in the UK Biobank

We applied RHE-mc to estimate genome-wide SNP heritability for 22 complex traits (6 quantitative and 16 binary traits) measured in the UK Biobank. We analyzed *N* = 291, 273 unrelated white British individuals and *M* = 459, 792 SNPs genotyped on the UK Biobank Axiom array (Methods). We ran RHE-mc with *B* = 10 and with SNPs divided into eight bins based on two MAF bins (0.01 ≤MAF< 0.05, MAF ≥ 0.05) and quartiles of the LD-scores. We compared the estimates from RHE-mc to those from LDSC, S-LDSC, SumHer, and GRE. Restricting our analysis to 18 traits for which the point estimate of genome-wide SNP heritability from RHE-mc is > 0.05, the estimates from S-LDSC, GRE, SumHer and LDSC were on average 2.5%, 10%, 25%, and 67% higher than RHE-mc (Figure 4). Relative to the simulation results, the estimates from S-LDSC are generally consistent with those from RHE-mc. This is likely due to the fact that, in simulations, our application of S-LDSC used only MAF bins. On the other hand, in real data, we used S-LDSC with the recommended baseline-LD annotations (including functional annotations).

**Figure 4:**
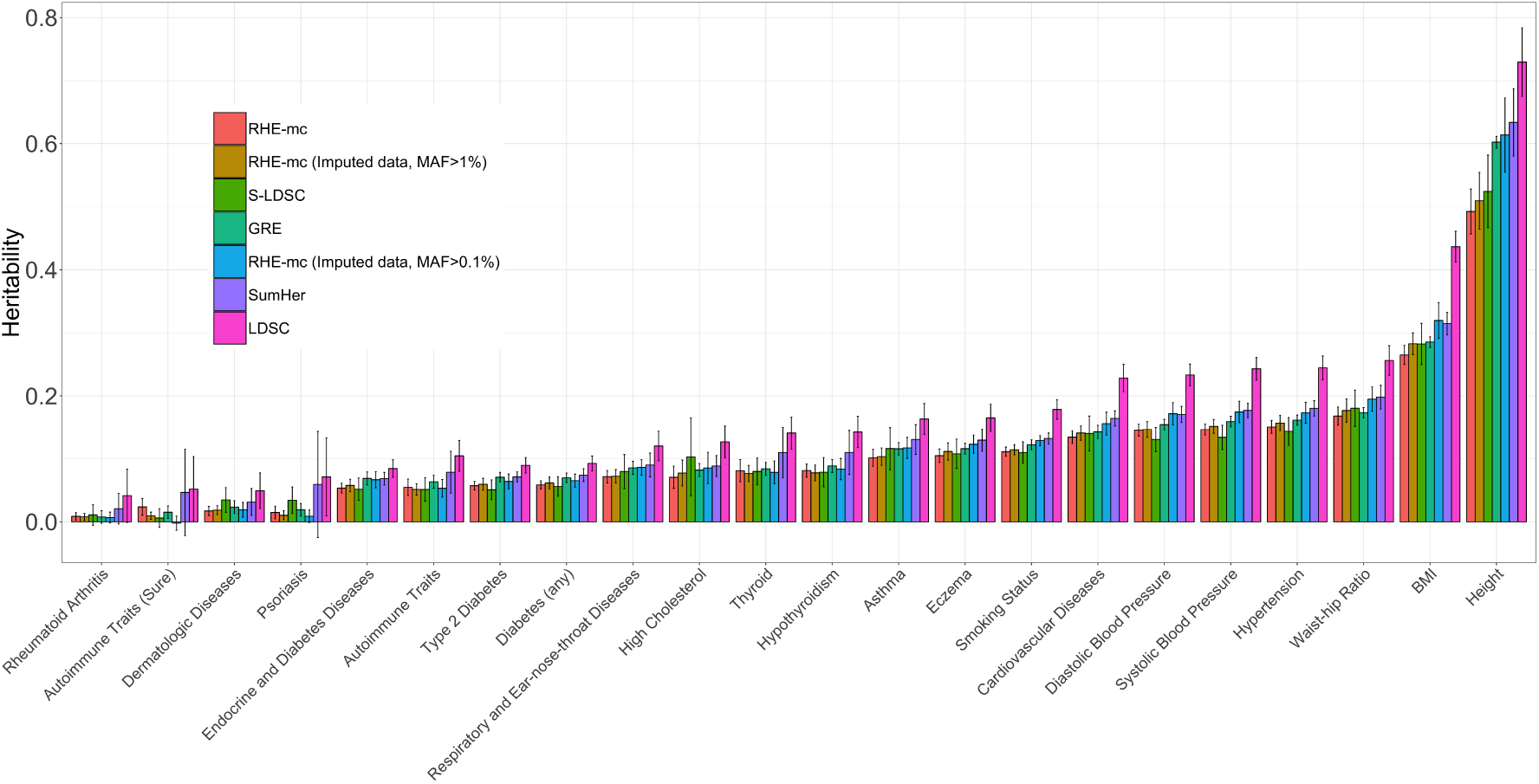
Estimates of genome-wide SNP heritability from RHE-mc, LDSC, S-LDSC, GRE, and SumHer for 22 complex traits and diseases in the UK Biobank. We restricted our analysis to *N* = 291, 273 unrelated white British individuals. We applied all methods to *M* = 459, 792 array SNPs (MAF> 1%). We ran S-LDSC with baseline-LD model. For every method, LD scores or LDAK weights are computed using in-sample LD among the SNPs, and we aim to estimate the SNP-heritability explained by the same set of SNPs. RHE-mc was applied to array SNPs with 8 MAF/LD bins. We also report RHE-mc estimates of genome-wide SNP heritability on *M* = 4, 824, 392 imputed SNPs (MAF > 1%) with 8 MAF/LD bins and *M* = 7, 774, 235 imputed SNPs (MAF > 0.1%) with 144 MAF/LD bins (see Methods). Black bars mark ±2 standard errors.

We then applied RHE-mc to estimate genome-wide heritability attributable to imputed variants. The genome-wide estimates of SNP heritability from RHE-mc on imputed SNPs (MAF> 1%) are concordant with the estimates from array SNPs (2.8% higher on average). We then analyzed *M* = 7, 774, 235 imputed genotypes with MAF > 0.1% using 144 bins formed by 4 LD bins and 36 MAF bins (Methods). Genome-wide SNP heritability estimates from RHE-mc on imputed SNPs (MAF> 0.1%) are 11.4% higher than RHE-mc on imputed SNPs (MAF> 1%). (see Figure 4, Supplementary Figure S7). 22

### Partitioning SNP heritability across allele frequency and LD bins

We used RHE-mc to partition SNP heritability of 22 complex traits across MAF and LD bins. We analysed *M* = 7, 774, 235 imputed SNPs with MAF > 0.1%. We used 144 bins formed by 4 LD bins and 36 MAF bins (see Methods). We compute the allelic effect size of SNPs in bin *k* as 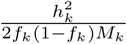 where 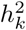 is the heritability estimated in bin *k, f*_*k*_ is the mean MAF in bin *k*, and *M*_*k*_ is the number of SNPs in bin *k*. We observe that allelic effect size increases with lower MAF and LD. For height, in the lowest quartile of LD scores, SNPs with MAF ≈ 0.1% have allelic effect sizes ≈ 27*x* ± 8 larger than SNPs with MAF ≈ 50%. Similarly, among SNPs with MAF ≈ 50%, SNPs in the lowest quartile of LD scores have allelic effect sizes ≈ 5*x* ± 1 larger than SNPs in the highest quartile (Figure 5 for height; other traits in Supplementary Figure S11). While these trends have been observed in previous studies [29, 9, 30], the ability of RHE-mc to jointly fit multiple variance components allows us to estimate effect sizes at SNPs with MAF as low as 0.1%. We caution that heritability estimates in bins of lowest MAF and high LD score tend to negative likely due to the low number of SNPs in this bin (we did not constrain our variance components estimates to be non-negative).

**Figure 5:**
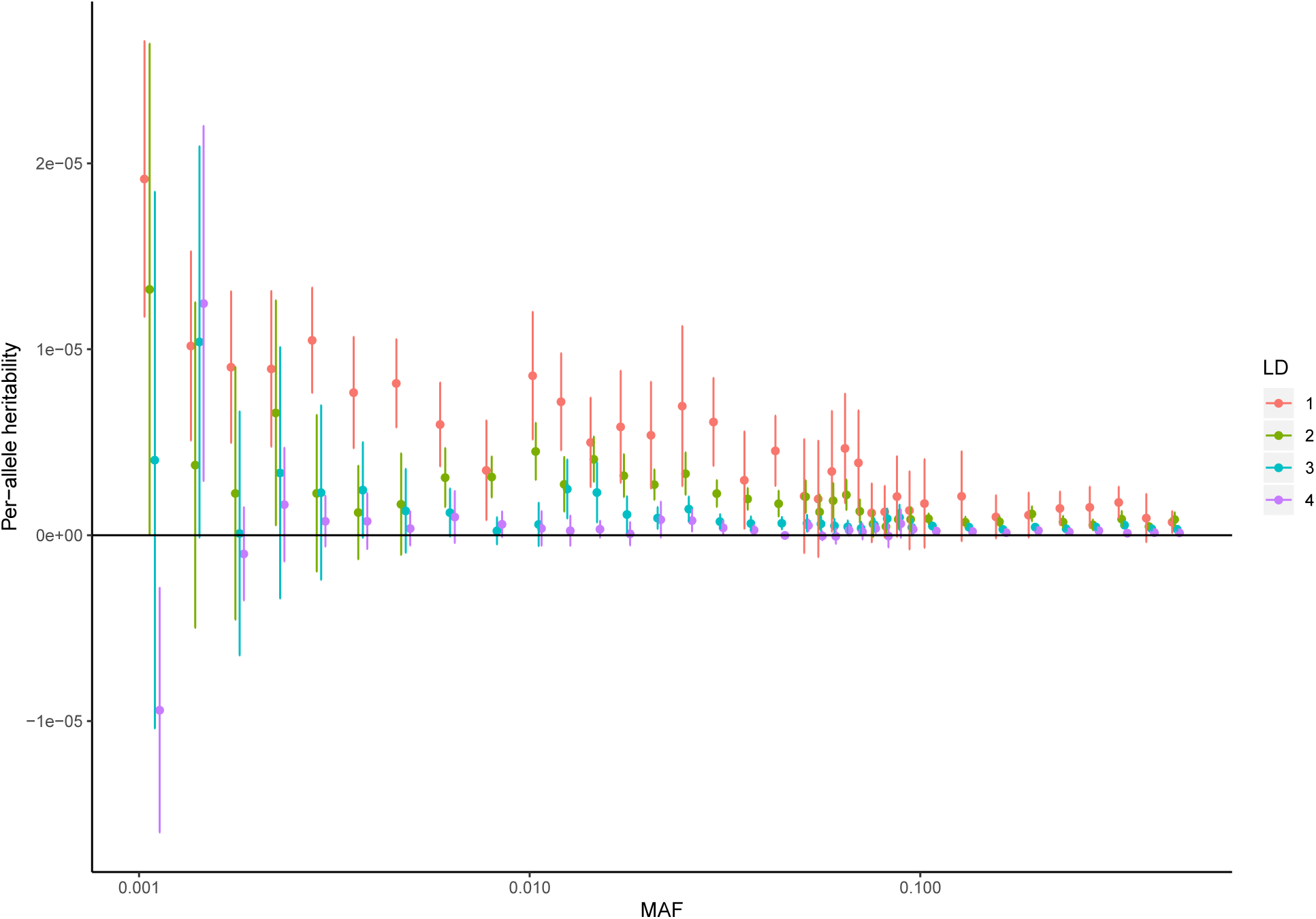
Per-allele heritability of height as a function of MAF: We applied RHE-mc to *N* = 291, 273 unrelated white British individuals and *M* = 7, 774, 235 imputed SNPs. SNPs were partitioned into 144 bins based on LD score (4 bins based on quartiles of the LD score with *i* denoting the *i*^*th*^ quartile) and MAF (36 MAF bins) (see Methods). Per allele heritability for bin *k* is defined as 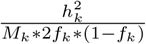 where 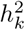 is the heritability attributed to bin *k, M*_*k*_ is the number of SNPs in bin *k*, and *f*_*k*_ is the average MAF in bin *k*. Bars mark ±2 standard errors. See Supplementary Figure S11 for results on all 22 traits.

### Partitioning heritability by functional annotations

The ability of RHE-mc to estimate variance components associated with a large number of overlapping annotations enables us to explore the contribution a variety of functional genomic annotations to trait heritability using individual-level data in the UK Biobank. We applied RHE-mc to jointly partition heritability of 22 complex traits across 28 functional annotations as defined in [7]. We restricted our analysis to *N* = 291, 273 unrelated white British individuals and *M* = 5, 670, 959 imputed SNPs (we restrict to SNPs with MAF > 0.1% which are also present in 1000 Genomes Project). We grouped the traits into five categories (autoimmune, diabetes, respiratory, anthropometric, cardiovascular); for a representative trait from each category, we report enrichment of each of the 28 functional annotations in Figure 6 (see Methods; for all traits see Supplementary Figure S10). Our results are largely concordant with previous studies [7, 9]: we observe enrichment of heritability across traits in conserved regions (Z-score > 3 in 15 traits). We also observe enrichment of heritability at FANTOM5 enhancers (labeled Enhancer Andersson in Figure 6) in asthma, eczema, autoimmune disorders (broad), hypothyroidism, and thyroid disorders (Z-score > 3) even though these annotations cover only 0.4% of the analyzed SNPs.

**Figure 6:**
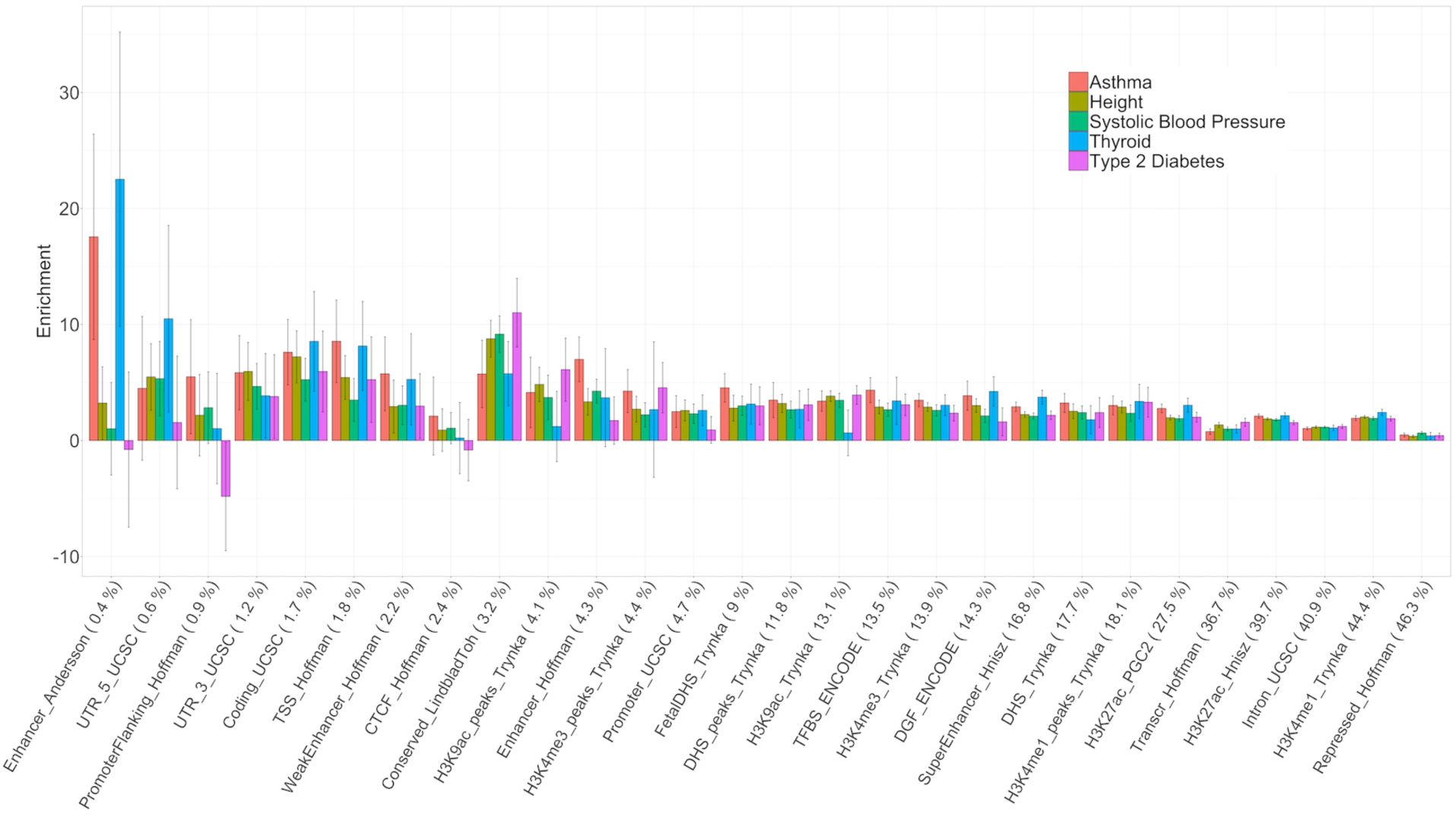
Enrichment of heritability across 28 functional annotations: We applied RHE-mc to *N* = 291, 273 unrelated white British individuals and *M* = 5, 670, 959 imputed SNPs (MAF > 0.1% and present in 1000 Genomes Project). SNPs were partitioned based on 28 functional annotations that were defined in a previous study [7]. We grouped 22 traits in the UK Biobank into five categories (autoimmune, diabetes, respiratory, anthropometric, cardiovascular). Here we plot enrichment of five traits (one representative trait per category). Black bars mark ± 2 standard errors. Annotations are ordered by the proportions of SNPs in that annotation (given in parentheses). See Supplementary Figure S10 for results on all 22 traits.

## Discussion

We have presented RHE-mc, an algorithm that can efficiently estimate multiple variance components on large-scale genoytpe data. In light of increasing evidence for SNP effect sizes that vary as a function of covariates such as MAF and LD and the bias associated with methods that fit only a single variance component [8], the ability to define flexible models endowed with multiple variance components is important to obtain unbiased estimates of fundamental quantities such as SNP heritability. We confirm that RHE-mc yields accurate genome-wide SNP heritability estimates under diverse genetic architectures. In applications to 22 complex traits in the UK Biobank, RHE-mc yields heritability estimates on array SNPs that are lower on average relative to S-LDSC and SumHer. We have explored the utility of RHE-mc in heritability partitioning analyses. These analyses show that allelic effect sizes tend to increase with a decrease in MAF and LD consistent with previous studies [9]. We also partitioned heritability across functional annotations to reveal enrichment of heritability at FANTOM5 enhancers in specific traits such as asthma and eczema.

We discuss several limitations of RHE-mc as well as directions for future work. First, the method-of-moments estimator underlying RHE-mc tends to yield slightly larger standard errors, on average, relative to REML estimators. The relative performance of the two methods likely depends on a number of aspects of the study design such as sample size, number of SNPs, the LD structure, relatedness patterns, and the underlying genetic architecture. Nevertheless, our method is designed to be applicable to massive datasets for which the heritability estimates are relatively precise. Developing scalable variance components estimators that are as efficient as REML-based methods is an important direction for future work. Second, this work has primarily explored the partitioning of heritability across discrete annotations. While we have shown how the methodology can be extended to continuous-valued annotations (Methods and Supplementary Information S8), it would be of interest to explore variation in trait heritability as a function of the value of an annotation. On the other hand, the ability of RHE-mc to fit many annotations allows the annotation to be divided into a sufficiently large number of bins. Third, we have applied RHE-mc to binary traits available in the UK Biobank treating these traits as continuous. Methods that explicitly model binary traits as well as the underlying ascertainment involved in case-control studies are likely to lead to more accurate heritability estimates [23, 31]. For example, the PCGC method [23] is an extension of HE regression and it would be of interest to develop a scalable randomized PCGC estimator. Fourth, RHE-mc requires access to individual-level genotype and phenotype data. Methods that only require summary statistic data (GRE [10], LDSC [24], and SumHer [25]) have the advantage of being applicable to datasets where acquiring access to individual-level data can be challenging [10]. Finally, our method could potentially lead to improvements in association testing, trait prediction, and understanding of polygenic selection.

## URLs

RHE-mc software: https://github.com/sriramlab/RHE-mc

## Acknowledgments

This research was conducted using the UK Biobank Resource under applications 33127 and 33297. We thank the participants of UK Biobank for making this work possible. We thank Rob Brown, Steven Gazal, and members of the Sankararaman and Pasaniuc labs for feedback on this manuscript. This work was funded by NIH grants R01HG009120 (B.P. and K.S.B.), R35GM125055 (S.S.), an Alfred P. Sloan Research Fellowship (S.S.), and a NSF grant III-1705121 (Y.W. and S.S.).

## Methods

### Multi-component Linear Mixed Model

RHE-mc attempts to fit the following variance components model:

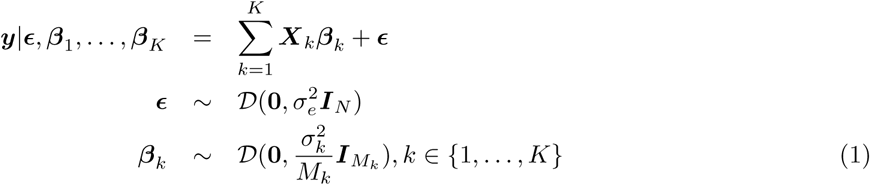

Here ***y*** is a *N*-vector of centered phenotypes. Here each of the *M* SNPs is assigned to one of *K* non-overlapping categories. Each category *k* contains *M*_*k*_ SNPs, *k* ∈ {1, …, *K*}, Σ_*k*_ *M*_*k*_ = *M*. Let ***X***_*k*_ be a *N* × *M*_*k*_ matrix where *x*_*k,n,m*_ denotes the standardized genotype for individual *n* at SNP *m* in category *k*. We have Σ_*n*_ *x*_*k,n,m*_ = 0 and 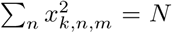 for *m* ∈ {1, 2, …, *M*_*k*_}. Let ***β***_*k*_ be a *M*_*k*_-vector of SNP effect sizes for the *k*-th category. In the above model, 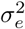 is the residual variance, and 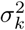 is the variance component of the *k*-th category. In this model, the total SNP heritability is defined as :

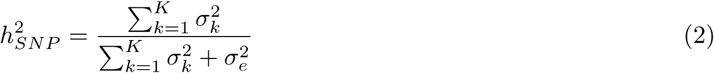

The SNP heritability of category *k* is defined as:

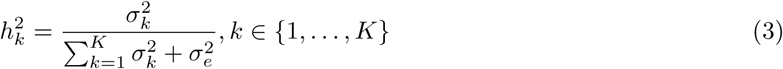

Enrichment in bin *k* is defined as:

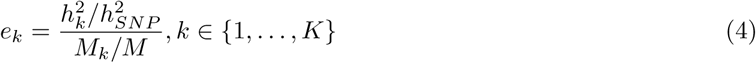

### Method-of-moments for estimating multiple variance components

To estimate the variance components, RHE-mc uses a Method-of-Moments (MoM) estimator that searches for parameter values so that the population moments are close to the sample moments [32]. Since 𝔼 [***y***] = 0, we derived the MoM estimates by equating the population covariance to the empirical covariance. The population covariance is given by:

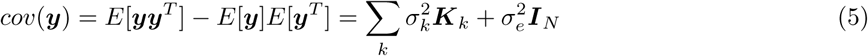

Here 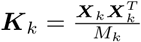 is the genetic relatedness matrix (GRM) computed from all SNPs of *k*-th category. Using ***yy***^T^ as our estimate of the empirical covariance, we need to solve the following least squares problem to find the variance components.

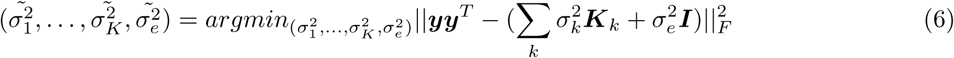

The MoM estimator satisfies the following normal equations:

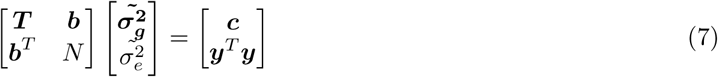

Here 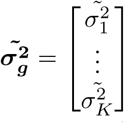, ***T*** is a *K* × *K* matrix with entries *T* = *tr*(***K***_*k*_ ***K***_*l*_), *k, l* ∈ {1, …, *K*}, ***b*** is a *K*-vector with entries *b*_*k*_ = *tr*(***K***_*k*_) = *N* (because ***X***_*k*_s is standardized), and ***c*** is a *K*-vector with entries *c*_*k*_ = ***y***^*T*^ ***K*_*k*_*y***. Each GRM ***K***_*k*_ can be computed in time 𝒪 (*N* ^2^*M*_*k*_) and 𝒪 (*N* ^2^) memory. Given *K* GRMs, the quantities *T*_*k,l*_, *c*_*k*_, *k, l* ∈ {1, …, *K*}, can be computed in 𝒪 (*K*^2^*N* ^2^). Given the quantities *T*_*k,l*_, *c*_*k*_, the normal equation 7 can be solved in 𝒪 (*K*^3^). Therefore, the total time complexity for estimating the variance components is 𝒪 (*N* ^2^*M* + *K*^2^*N* ^2^ + *K*^3^).

### RHE-mc: Randomized estimator of multiple variance components

The key bottleneck in solving the normal equation 7 is the computation of *T*_*k,l*_, *k, l* ∈ {1, …, *K*} which takes 𝒪 (*N* ^2^*M*). Instead of computing the exact value of *T*_*k,l*_, we use an unbiased estimator of the trace [33] based on the following identity: for a given *N* × *N* matrix ***C, z***^*T*^ ***Cz*** is an unbiased estimator of *tr*(***C***) (*E*[***z***^*T*^ ***Cz***] = *tr*[***C***]) where ***z*** be a random vector with mean zero and covariance ***I***_*N*_. Hence, we can estimate the values *T*_*k,l*_, *k, l* ∈ {1, …, *K*} as follows:

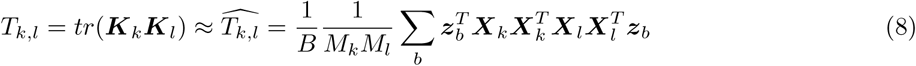

Here ***z***_1_, …, ***z***_*B*_ are *B* independent random vectors with zero mean and covariance ***I***_*N*_. We draw these random vectors independently from a standard normal distribution. Computing *T*_*k,l*_ using the unbiased estimator involves four multiplications of sub-matrices of the genotype matrix with a vector, repeated *B* times. Therefore, the total running time for estimating the value *T*_*k,l*_ is 𝒪 (*NMB*).

Moreover, we can leverage the structure of the genotype matrix which only contains entries in {0, 1, 2}. For a fixed genotype matrix ***X***_*k*_, we can improve the per iteration time complexity of matrix-vector multiplication from 𝒪 (*NM*) to 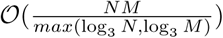 by using the Mailman algorithm [34]. Solving the normal equations takes 𝒪 (*K*^3^) time so that the overall time complexity of our algorithm is 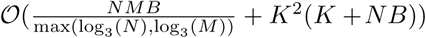.

RHE-mc uses a block Jackknife to estimate standard errors. In Supplementary Information S1, we show how the block Jackknife estimates can be computed with little additional computational overhead. Further, we also show how covariates can be efficiently included in the model (Supplementary Information S2).

### Multi-component LMM with overlapping annotations

RHE-mc can also be applied in the setting where annotations overlap. Following [7], the heritability of SNPs belong to annotation *k* is defined as:

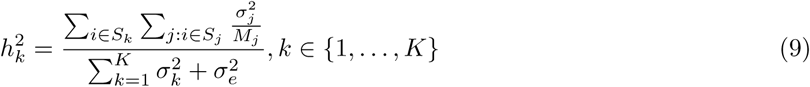

where *S*_*k*_ is the set of SNPs in *k*-th annotation and *M*_*k*_ = |*S*_*k*_ |. Enrichment in bin *k* is defined as 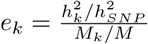.

### Multi-component LMM with continuous annotations

We have described the derivation of RHE-mc using binary annotations. Here we extend the method to continuous-value annotation as follows :

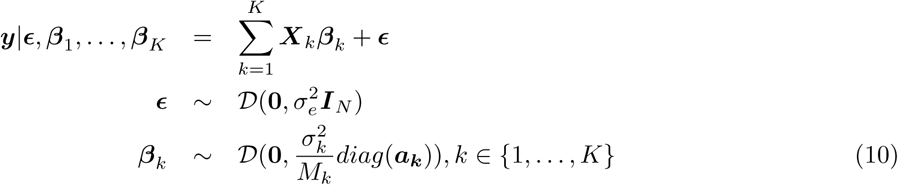

This model is totally similar to the model 10 except that here we assume that the variances of effect sizes depend on continuous-valued annotation. Let ***a***_*k*_ be a *M*_*k*_-vector where ***a***_*k*,***i***_ is the value of *k*-th annotation at SNP *i* (the elements of ***a***_*k*_ must be non-negative). Let *S*_*k*_ be the set of SNPs belong to annotation *k*. In this model, the SNP heritability of annotation k is defined as :

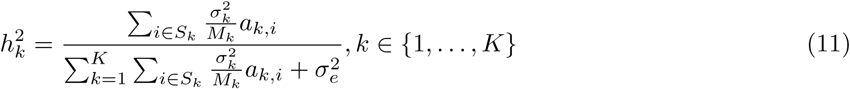

To estimate the variance components of this new model, we only need to replace ***X***_*k*_ with 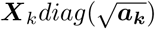 in the equation 5 for every annotation *k*. We assessed the accuracy of RHE-mc in estimating variance components with continuous annotation in Supplementary Information S8.

### Simulations

We performed simulations to compare the performance of RHE-mc with several state-of-the-art methods for heritability estimation that cover the spectrum of methods that have been proposed.

We considered two simulation settings. In the large-scale simulation setting, we simulated phenotypes for the full set of UK Biobank genotypes consisting of *M* = 593, 300 array SNPs and *N* = 337, 205 individuals. We obtained the individuals by keeping unrelated white British individuals which are > 3^*rd*^ degree relatives (defined as pairs of individuals with kinship coefficient < 1*/*2^(9*/*2)^)[17], and removing individuals with putative sex chromosome aneuploidy. The small-scale setting was designed so that we could compare the accuracies of RHE-mc to REML methods. In this setting, we simulated phenotypes from a subsampled set of genotypes from the UK Biobank data genotypes used in large scale simulation [35]. Specifically, we chose randomly a subset of *N* = 10, 000 individuals from the large scale data. Therefore, in small scale, we have *M* = 593, 300 array SNPs and *N* = 10, 000 individuals. We simulated phenotypes from genotypes using the following model which is used in [10, 8]:

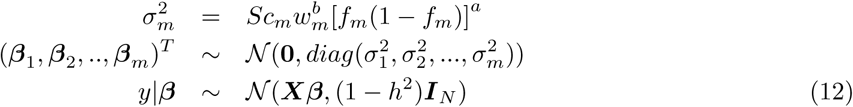

where *S* is a normalizing constant chosen so that 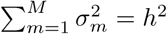. Here *h*^2^ ∈ [0, 1], *a* ∈ {0, 0.75}, *b* ∈ {0, 1}, *β*_*m*_, *f*_*m*_ and *w*_*m*_ are the effect size, the minor allele frequency and LDAK score of *m*^*th*^ SNP respectively. Let *c*_*m*_ ∈ {0, 1} be an indicator variable for the causal status of SNP *m*. The LD score of a SNP is defined to be the sum of the squared correlation of the SNP with all other SNPs that lie within a specific distance, and the LDAK score of a SNP is computed based on local levels of LD such that the LDAK score tends to be higher for SNPs in regions of low LD [36]. The above models relating genotype to phenotype are commonly used in methods for estimating SNP heritability: the GCTA Model (when *a* = *b* = 0 in Equation 12), which is used by the software GCTA [27] and LD Score regression (LDSC) [24], and the LDAK Model (where *a* = 0.75, *b* = 1 in Equation 12) used by software LDAK [36]. Moreover, under each model, we varied the proportion and minor allele frequency (MAF) of causal variants (CVs). Proportion of causal variants were set to be either 100% or 1%, and MAF of causal variants drawn uniformly from [0, 0.5] or [0.01, 0.05] or [0.05, 0.5] to consider genetic architectures that are either infinitesimal or sparse as well genetic architectures that include a mixture of common and rare SNPs as well as one that includes only common SNPs. The true heritability were chosen from {0.1, 0.25, 0.5, 0.8}

We generated 100 sets of simulated phenotypes for each setting of parameters and report accuracies averaged over these 100 sets.

### Comparisons

For the large-scale simulations, we compared RHE-mc to methods that rely on summary statistics for estimating heritability. Among the summary statistic methods, LD score regression (LDSC) [24] uses the slope from the GWAS *χ*^2^ statistics regressed on the LD scores to estimate heritability. Stratified LD score regression (S-LDSC) [7] is an extension of LDSC for partitioning heritability from summary statistics. SumHer is the summary statistic analog of LDAK [25]. We ran S-LDSC with 10 binary MAF bin annotations defined such that each bin contains exactly 10% of the typed SNPs; this is intended to mirror the 10 MAF bin annotations in the S-LDSC “baseline-LD model” [29] (see Supplementary Table S5). To run SumHer, we used the LDAK software to compute the default “LDAK weights” using in-sample LD [36, 37, 25]. We then computed “LD tagging” using 1-Mb windows centered on each SNP as recommended [25]. To do a fair comparison we computed LD scores for LDSC, S-LDSC, GRE, and SumHer by using in-sample LD among the M SNPs, and in all simulations we aim to estimate the SNP-heritability explained by the same set of M SNP. We described the parameter settings of summary statistic methods in Supplementary Information S9.

For the small-scale simulations, we compared RHE-mc to GCTA-mc and HE-mc [27]. GCTA-mc and HE-mc are the extensions of GCTA and HE to a multi-component LMM respectively where the variance components are typically defined by binning SNPs according to their MAF as well as local LD [8]. We ran both GCTA-mc and RHE-mc using 24 bins formed by the combination of 6 bins based on MAF (MAF 0.01,0.01 <MAF ≤ 0.02,0.02 <MAF ≤ 0.03,0.03 <MAF ≤ 0.4,0.04 <MAF ≤ 0.05,MAF> 0.05) as well as 4 bins based on quartiles of the LDAK score of a SNP. We ran both GCTA-mc and RHE-mc allowing for estimates of a variance component to be negative.

For comparisons of runtime, we compared RHE-mc to GCTA [27] and BOLT-REML [4] which is a computationally efficient approximate method to compute the REML estimator. We ran all methods with 22 components (one for each chromosome). We also ran RHE-mc with ≈ 300 components (corresponding to 10 Mb bins) on the UK Biobank genotype (Supplementary Figure S8). To create our largest dataset, we replicate individuals from the UK Biobank and a subset of the imputed SNPs to obtain a dataset with one million individuals and SNPs. We use the latest versions of BOLT-REML (Version 2.3.2) and GCTA (Version 1.92.1) in our comparison. All comparisons are performed on an Intel(R) Xeon(R) CPU 2.10 GHz server with 128 GB RAM.

### Heritability estimates in the UK Biobank

We estimated SNP-heritability for 22 real complex traits (6 quantitative, 16 binary) in the UK Biobank [17]. In this study, we restricted our analysis to SNPs that were present in the UK Biobank Axiom array used to genotype the UK Biobank. SNPs with greater than 1% missingness and minor allele frequency smaller than 1% were removed. Moreover, SNPs that fail the Hardy-Weinberg test at significance threshold 10^−7^ were removed. We restricted our study to self-reported British white ancestry individuals which are > 3^*rd*^ degree relatives that is defined as pairs of individuals with kinship coefficient < 1*/*2^(9*/*2)^ [17]. Furthermore, we removed individuals who are outliers for genotype heterozygosity and/or missingness. Finally we obtained a set of *N* = 291, 273 individuals and *M* = 459, 792 SNPs to use in the real data analyses. We included age, sex, and the top 20 genetic principal components (PCs) as covariates in our analysis for all traits. We used PCs precomputed by the UK Biobank from a superset of 488, 295 individuals. Additional covariates were used for waist-to-hip ratio (adjusted for BMI) and diastolic/systolic blood pressure (adjusted for cholesterol-lowering medication, blood pressure medication, insulin, hormone replacement therapy, and oral contraceptives).

### Heritability partitioning

In our initial analysis, we removed SNPs with greater than 1% missingness and minor allele frequency smaller than 1%. Moreover, we removed SNPs that fail the Hardy-Weinberg test at significance threshold 10^−7^ as well as SNPs that lie within the MHC region (Chr6: 25–35 Mb) to obtain 4, 824, 392 SNPs. We restricted our study to individuals self-reported British white ancestry individuals which are > 3^*rd*^ degree relatives that is defined as pairs of individuals with kinship coefficient < 1*/*2^(9*/*2)^ [17]. Furthermore, we removed individuals who are outliers for genotype heterozygosity and/or missingness. Finally, we obtained 291, 273 individuals. We partitioned SNPs into eight bins based on two MAF bins (MAF ≤ 0.05, MAF> 0.05) and quartiles of the LD-scores. For each bin *k*, we computed the heritability enrichment as the ratio of the percentage of heritability explained by SNPs in bin *k* to the the percentage of SNPs in bin *k*.

We considered an additional analysis in which we included SNPs with MAF > 0.1% resulting in *N* = 291, 273 unrelated white British individuals and *M* = 7, 774, 235 imputed SNPs (MAF > 0.1%). We defined 144 bins based on 4 LD bins and 36 MAF bins. The four LD bins are defined based on quartile of LD-scores, and 36 MAF bins are defined based on 9-quantile of the following four intervals: 0.001 ≤MAF≤ 0.01,, 0.01 <MAF≤ 0.05, 0.05 ≤MAF≤ 0.10, 0.10 <MAF≤ 0.50.

## Data availability

Access to the UK Biobank resource is available via application at: http://www.ukbiobank.ac.uk.

## Code availability

RHE-mc software is open-source software freely available at: https://github.com/sriramlab/RHE-mc

