## Supplementary information for "Efficient variance components analysis across millions of genomes"

### Supplementary Information: Efficient variance components analysis across millions of genomes

#### Contents

**S1 Computing the Standard Errors of the RHE-mc estimates** **2**

**S2 Including covariates in RHE-mc** **2**

**S3 Implementation details** **3**
S3.1 Streaming version . . . . . . . . . . . . . . . . . . . . . . . . . . . . . . . . . . . . . . . 3

**S4 Jackknife standard errors** **5**

**S5 Number of random vectors** **5**

**S6 Simulated data** **7**

**S7 Parameter settings for summary statistics methods** **13**

**S8 Continuous annotations** **13**

**S9 Power as a function of annotation size** **14**

**S10 Array UK Biobank data** **21**

### S1 Computing the Standard Errors of the RHE-mc estimates

We obtain standard errors for RHE-mc using a block jackknife [1]. A jackknife subsample is created by leaving out a subset of observations from a dataset. The jackknife estimate of a parameter can be found by estimating the parameter for each subsample omitting the $i$-th jackknife block. A naive way to compute jackknife estimate requires computing the estimator of the parameters for every subsample. For instance, in our problem, if we define $J$ jackknife blocks, then we need to run RHE-mc for every subsample which takes $\mathcal{O}(J(\frac{NMB}{\max(\log_3(N),\log_3(M))} + K^2(K+NB)))$. We propose an efficient way to compute the jackknife estimate in time $\mathcal{O}(\frac{NMB}{\max(\log_3(N),\log_3(M))} + JK^2(K+NB))$.

Let $\boldsymbol{X}$ be a $N \times M$ matrix of standardized genotypes where $N$ and $M$ are the number of individuals and SNPs respectively. To generate $J$ jackknife subsamples, we partition $X$ into $J$ non-overlapping blocks $\boldsymbol{X}^{(1)}, \ldots, \boldsymbol{X}^{(J)}$ such that $\boldsymbol{X} = [\boldsymbol{X}^{(1)}, \boldsymbol{X}^{(2)}, \ldots, \boldsymbol{X}^{(J)}]$ . Note that for every $j$, $\boldsymbol{X}^{(j)}$ is a $N \times M_j$ matrix where $M_j$ is the number of SNPs in the $j$-th block.

We create the $j$-th jackknife subsample by removing the $j$-th block $\boldsymbol{X}^{(j)}$ from $\boldsymbol{X}$. To estimate the variance components of the $j$-th jackknife subsample, we need to compute the corresponding quantities of the $j$th subsample in the normal equations (Methods). Let $\boldsymbol{K}_k^{(-j)}$ be the GRM of the $k$-th partition which is created by removing the $j$-th block $\boldsymbol{X}^{(j)}$ from $\boldsymbol{X}$ where $k \in \{1, \ldots, K\}$, $j \in \{1, \ldots, J\}$. In Algorithm S3.1, we show how we can compute $tr(\widehat{\boldsymbol{K}_k^{(-j)}\boldsymbol{K}_l^{(-j)}})$ and $\boldsymbol{y}^T\boldsymbol{K}_i^{(-j)}\boldsymbol{y}$, for all $k, l \in \{1, \ldots, K\}$, $j \in \{1, \ldots, J\}$ efficiently.

### S2 Including covariates in RHE-mc

We can extend the LMM to include covariates as follows:

$$\boldsymbol{y}|\boldsymbol{\epsilon}, \boldsymbol{\beta}_1, \ldots, \boldsymbol{\beta}_k = \boldsymbol{W}\boldsymbol{\alpha} + \sum_k \boldsymbol{X}_k\boldsymbol{\beta}_k + \boldsymbol{\epsilon} \tag{1}$$

Here $\boldsymbol{W}$ is a $N \times C$ matrix of covariates while $\boldsymbol{\alpha}$ is a $C$-vector of fixed effects.

It is easy to see that the matrix $\boldsymbol{V} = \boldsymbol{I}_N - \boldsymbol{W}(\boldsymbol{W}^T\boldsymbol{W})^{-1}\boldsymbol{W}^T$ is symmetric and idempotent ($\boldsymbol{V}^2 = \boldsymbol{V}$) of rank $N - C$. Therefore, we consider the eigendecomposition of $\boldsymbol{V} = \boldsymbol{E}\boldsymbol{D}\boldsymbol{E}^T$, where $\boldsymbol{D}$ is a diagonal matrix with $N - C$ ones and $C$ zeros on the diagonal (we can assume that first $N - C$ elements are one). Now let the matrix $\boldsymbol{U}_{N\times(N-C)}$ represent the first $N - C$ columns of $\boldsymbol{E}$. It is not hard to see that $\boldsymbol{U}$ satisfies $\boldsymbol{U}^T\boldsymbol{U} = \boldsymbol{I}_{N-C}$, $\boldsymbol{U}\boldsymbol{U}^T = \boldsymbol{V}$, $\boldsymbol{U}^T\boldsymbol{W} = 0$. Now we multiplying by $\boldsymbol{U}^T$ on both sides of the above equation:

$$\boldsymbol{U}^T\boldsymbol{y} = \boldsymbol{U}^T\sum_k \boldsymbol{X}_k\boldsymbol{\beta}_k + \boldsymbol{U}^T\boldsymbol{\epsilon} \tag{2}$$

$$cov(\boldsymbol{U}^T\boldsymbol{y}) = E[\boldsymbol{U}^T\boldsymbol{y}(\boldsymbol{U}^T\boldsymbol{y})^T] - E[\boldsymbol{U}^T\boldsymbol{y}]E[\boldsymbol{U}^T\boldsymbol{y}] \tag{3}$$

The matrix $\boldsymbol{U}^T$ is constant and the vector $\boldsymbol{y}$ is random. Therefore, we have $E[\boldsymbol{U}^T\boldsymbol{y}] = \boldsymbol{U}^TE[\boldsymbol{y}]$.

$$\boldsymbol{U}^T\boldsymbol{y}(\boldsymbol{U}^T\boldsymbol{y})^T = (\boldsymbol{U}^T\sum_k \boldsymbol{X}_k\boldsymbol{\beta}_k + \boldsymbol{U}^T\boldsymbol{\epsilon})(\boldsymbol{U}^T\sum_k \boldsymbol{X}_k\boldsymbol{\beta}_k + \boldsymbol{U}^T\boldsymbol{\epsilon})^T = \tag{4}$$

$$\sum_i\sum_j \boldsymbol{U}^T\boldsymbol{X}_i\boldsymbol{\beta}_i(\boldsymbol{U}^T\boldsymbol{X}_j\boldsymbol{\beta}_j)^T + (\boldsymbol{U}^T\boldsymbol{\epsilon})\sum_i(\boldsymbol{U}^T\boldsymbol{X}_i\boldsymbol{\beta}_i)^T + \sum_i \boldsymbol{U}^T\boldsymbol{X}_i\boldsymbol{\beta}_i(\boldsymbol{U}^T\boldsymbol{\epsilon})^T + \boldsymbol{U}^T\boldsymbol{\epsilon}(\boldsymbol{U}^T\boldsymbol{\epsilon})^T$$

Hence

$$E[\boldsymbol{U}^T\boldsymbol{y}(\boldsymbol{U}^T\boldsymbol{y})^T] = \sum_k \frac{\sigma^2_{g_k}}{M_k}(\boldsymbol{U}^T\boldsymbol{X}_k)(\boldsymbol{U}^T\boldsymbol{X}_k)^T + \sigma^2_\epsilon \boldsymbol{U}^T\boldsymbol{U} \tag{5}$$

Using $\boldsymbol{K}_k = \frac{\boldsymbol{X}_k\boldsymbol{X}_k^T}{M_k}$, we have:

$$cov(\boldsymbol{U}^T\boldsymbol{y}) = \boldsymbol{U}^T(\sum_k \sigma^2_{g_k}\boldsymbol{K}_k)\boldsymbol{U} + \sigma^2_\epsilon \boldsymbol{I}_{N-C} \tag{6}$$

The MoM estimator is obtained by solving the following ordinary least squares problem:

$$(\tilde{\sigma_1^2}, \dots, \tilde{\sigma_K^2}, \tilde{\sigma_e^2}) = argmin_{(\sigma_1^2,\dots,\sigma_K^2,\sigma_e^2)} ||\boldsymbol{U}^T\boldsymbol{y}(\boldsymbol{U}^T\boldsymbol{y})^T - \boldsymbol{U}^T(\sum_k \sigma_k^2\boldsymbol{K}_k)\boldsymbol{U} - \sigma^2_\epsilon\boldsymbol{I}_{N-C}||_F^2 \tag{7}$$

We need to solve the following normal equations to estimate the variance components.

$$\begin{bmatrix} \boldsymbol{T} & \boldsymbol{b} \\ \boldsymbol{b}^T & N-C \end{bmatrix} \begin{bmatrix} \sigma_1^2 \\ \vdots \\ \sigma_k^2 \\ \sigma_e^2 \end{bmatrix} = \begin{bmatrix} \boldsymbol{c} \\ \boldsymbol{y}^T\boldsymbol{V}\boldsymbol{y} \end{bmatrix} \tag{8}$$

Here $\boldsymbol{V} = \boldsymbol{I}_N - \boldsymbol{W}(\boldsymbol{W}^T\boldsymbol{W})^{-1}\boldsymbol{W}^T$ and $\boldsymbol{T}$ is a $K \times K$ matrix where $T_{k,l} = tr(\boldsymbol{K}_k\boldsymbol{V}\boldsymbol{K}_l\boldsymbol{V})$, and $b$ is a $K-$vector where $b_k = tr(\boldsymbol{V}\boldsymbol{K}_k)$, and $\boldsymbol{c}$ is a $K$- vector where $c_k = \boldsymbol{y}^T\boldsymbol{V}\boldsymbol{K}_k\boldsymbol{V}\boldsymbol{y}$. Commonly, the number of covariates $C$ is small (tens to hundreds) so that including covariates does not significantly affect the computational cost. The cost of computing the elements of the normal equations 8 includes the cost of inverting $\boldsymbol{W}^T\boldsymbol{W}$ which is a $C \times C$ matrix and multiplying $\boldsymbol{W}$ by a real-valued $N$-vector which can be done in $\mathcal{O}(C^3 + NC)$.

### S3 Implementation details

#### S3.1 Streaming version

Here we describe the streaming version of RHE-mc algorithm. In Methods section, we showed that our MoM estimator satisfy the following normal equation.

$$\begin{bmatrix} \boldsymbol{T} & \boldsymbol{b} \\ \boldsymbol{b}^T & N \end{bmatrix} \begin{bmatrix} \tilde{\boldsymbol{\sigma_g^2}} \\ \tilde{\sigma_e^2} \end{bmatrix} = \begin{bmatrix} \boldsymbol{c} \\ \boldsymbol{y}^T\boldsymbol{y} \end{bmatrix} \tag{9}$$

Here $\tilde{\boldsymbol{\sigma_g^2}} = \begin{bmatrix} \tilde{\sigma_1^2} \\ \vdots \\ \tilde{\sigma_K^2} \end{bmatrix}$, $\boldsymbol{T}$ is a $K \times K$ matrix with entries $T_{k,l} = tr(\boldsymbol{K}_k\boldsymbol{K}_l), k, l \in \{1, \dots, K\}$, $\boldsymbol{b}$ is a $K$-vector with entries $b_k = tr(\boldsymbol{K}_k) = N$ (because $\boldsymbol{X}_k$s is standardized ), and $\boldsymbol{c}$ is a $K$-vector with entries $c_k = \boldsymbol{y}^T\boldsymbol{K}_k\boldsymbol{y}$. Here we estimate $T_{k,l}$ as follows :

$$T_{k,l} = tr(\boldsymbol{K}_k\boldsymbol{K}_l) \approx \widehat{T_{k,l}} = \frac{1}{B}\frac{1}{M_kM_l}\sum_b \boldsymbol{z}_b^T\boldsymbol{X}_k\boldsymbol{X}_k^T\boldsymbol{X}_l\boldsymbol{X}_l^T\boldsymbol{z}_b \tag{10}$$

Here $\boldsymbol{z}_1, \dots, \boldsymbol{z}_B$ are $B$ independent random vectors with zero mean and covariance $\boldsymbol{I}_N$.

We read genotype matrix $\boldsymbol{X}_k$ for every $k \in \{1, \ldots, K\}$ block by block. We define $J$ blocks over $\boldsymbol{X}_k$ by partitioning the columns of $\boldsymbol{X}_k$ to $J$ groups such that $\boldsymbol{X_k} = [\boldsymbol{X_k}^{(1)} \ldots \boldsymbol{X_k}^{(J)}]$.

**Algorithm 1:** Streaming version of RHE-mc

```
for every genotype matrix k ∈ {1,..,K} do
    for every block j ∈ {1,..,J} do
        Read X_k^j
        for every random vector b ∈ {1,..,B} do
            Z_(k,j,b) = X_k^j X_k^j^T z_b
        end
        v = X_k^j^T y
        H_(k,j) = v^T v
        Release the memory allocated to X_k^j
    end
end
Let U_{k,b,0} = Σ_j Z_(k,j,b), and
Let U_{k,b,j} = U_{k,b,0} − Z_(k,j,b), for every k, b, j.
Let V_{k,0} = Σ_j H_{k,j},
Let V_{k,j} = V_{k,0} − H_{k,j} for every k, j
for every j ∈ {0,1,..,J} do
    for every pairs of genotype matrices k and l ∈ {1,..,K} do
        T̂_{k,l} = (1/B) (1/(M_k M_l)) Σ_b U_(k,j,b)^T U_(l,j,b)
    end
    for every genotype matrix k ∈ {1,..,K} do
        c_k = (1/M_k) V_(k,j)^T V_(k,j)
    end
    Solve the normal equation for j^th sub-sample (j = 0 corresponds to the original genotype
      matrix used for computing the point estimates)
end
Compute the jackknife SE from the point estimates of J sub-samples.
```

In the above algorithm, the 3-D matrices $Z$ and $U$ need $\mathcal{O}(JKBN)$ memory, the 2-D matrices $V$ and $H$ need $\mathcal{O}(JN)$ memory. So the total space complexity will be $\mathcal{O}(JKBN)$. The total running time of this implementation is $\mathcal{O}(\frac{NMB}{\max(\log_3(N), \log_3(M))} + JK^2(K + NB)))$. For simplicity we assume that the streaming blocks are same as jackknife blocks. However, we can set the size of the streaming blocks to be different from the jackknife blocks to make the algorithm more efficient in terms of memory usage.

#### S4 Jackknife standard errors

| Genetic architecture | | | | True SE | Jackknife SE |
|---|---|---|---|---|---|
| Percentage of causal SNPs | MAF of causal SNPs | MAF and LD coupling | True $h^2$ | | |
| 0.01 | [0.01,0.05] | a=b=0 | 0.5 | 0.012 | 0.013 |
| 0.01 | [0.01,0.05] | a=0,b=1 | 0.5 | 0.018 | 0.015 |
| 0.01 | [0.01,0.05] | a=0.75,b=0 | 0.5 | 0.016 | 0.015 |
| 0.01 | [0.01,0.05] | a=0.75,b=1 | 0.5 | 0.013 | 0.013 |
| 0.01 | [0.0,0.5] | a=b=0 | 0.5 | 0.009 | 0.013 |
| 0.01 | [0.0,0.5] | a=0,b=1 | 0.5 | 0.016 | 0.014 |
| 0.01 | [0.0,0.5] | a=0.75,b=0 | 0.5 | 0.018 | 0.019 |
| 0.01 | [0.0,0.5] | a=0.75,b=1 | 0.5 | 0.012 | 0.015 |
| 0.01 | [0.05,0.5] | a=b=0 | 0.5 | 0.012 | 0.015 |
| 0.01 | [0.05,0.5] | a=0,b=1 | 0.5 | 0.021 | 0.017 |
| 0.01 | [0.05,0.5] | a=0.75,b=0 | 0.5 | 0.014 | 0.014 |
| 0.01 | [0.05,0.5] | a=0.75,b=1 | 0.5 | 0.015 | 0.017 |
| 1.0 | [0.0,0.5] | a=b=0 | 0.5 | 0.007 | 0.007 |
| 1.0 | [0.0,0.5] | a=0,b=1 | 0.5 | 0.007 | 0.007 |
| 1.0 | [0.0,0.5] | a=0.75,b=0 | 0.5 | 0.006 | 0.006 |
| 1.0 | [0.0,0.5] | a=0.75,b=1 | 0.5 | 0.007 | 0.008 |

Table S1: **Comparison of true SE with jackknife SE under 16 different genetic architectures**: We defined 100 blocks over SNPs to estimate block jackknife SE. We ran RHE-mc with 24 bins based on 6 MAF bins and 4 LDAK score bins (see Methods). True SE is computed from 100 replicates for every setting. Jackknife SE yields estimates of true SE with relative bias −3% on average over 16 genetic architectures.

#### S5 Number of random vectors

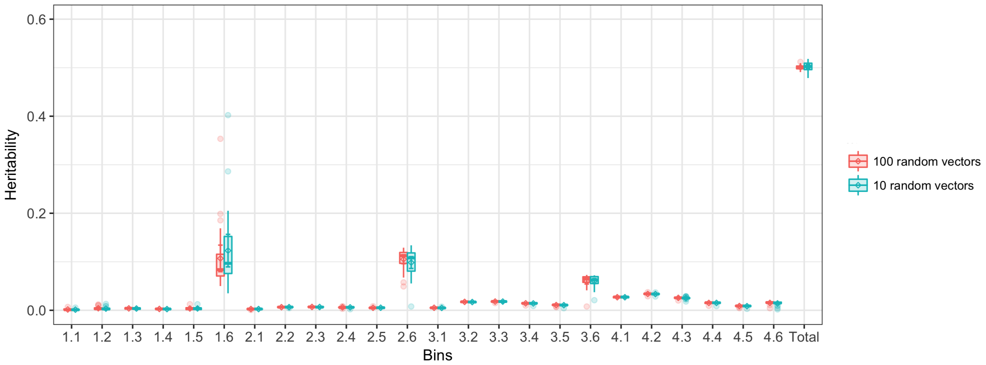

Figure S1: **Comparison of RHE-mc heritability estimates with $B = 10$ and $B = 100$ random vectors on large-scale simulated data (M=590K array SNPs and N=337K individuals)**: We ran RHE-mc with 24 bins( based on 6 MAF bins and 4 LDAK bins, see Methods). Standard errors are computed from 100 replicates. Here x-axis represents the bins ($i.j$ denotes the bin defined based on $i$-th ldak bin and $j$-th MAF bin) and y-axis represents the heritability.

| Number of random vectors | Point estimate (true SE) | Point estimate (SE of the estimator due to randomization) |
|---|---|---|
| 10 | 0.24 (0.06) | 0.24 (0.02) |
| 100 | 0.24 (0.05) | 0.25 (0.001) |

Table S2: Comparison of RHE-mc estimates with B=10 and B=100 on small scale (M=590K array SNPs and N=10k individuals). Here we quantify the contribution of randomization to the SE of the estimator. Here the true total heritability is 0.25. We first computed the SE of RHE-mc for $B = 10$ and $B = 100$ from 100 simulation replicates (second column). We then computed the SE of the estimates (due to the randomization) for a single replicate. For $B = 10$, randomization contributes about a third of the total SE ($\frac{0.02}{0.06}$).

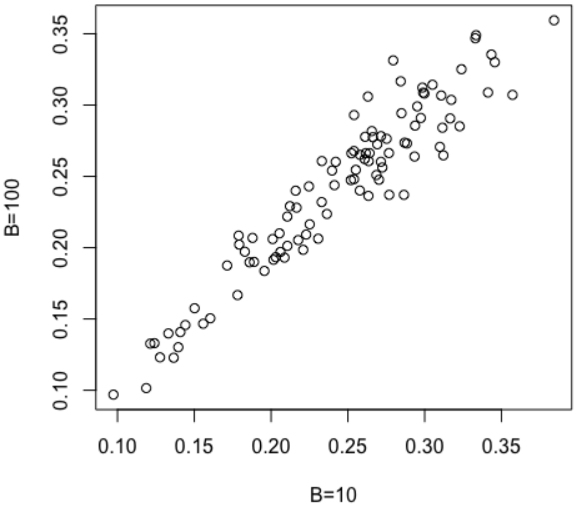

Figure S2: Comparison of RHE-mc estimates with B=10 and B=100 on small scale (M=590K array SNPs and N=10k individuals): We simulated 100 phenotpyes such that the true total heritability is 0.25. Here x-axis represents the RHE-mc estimates when $B = 10$, and y-axis represents RHE-mc estimates when $B = 100$.

### S6 Simulated data

#### S6.1 Heritability partitioning on small-scale data

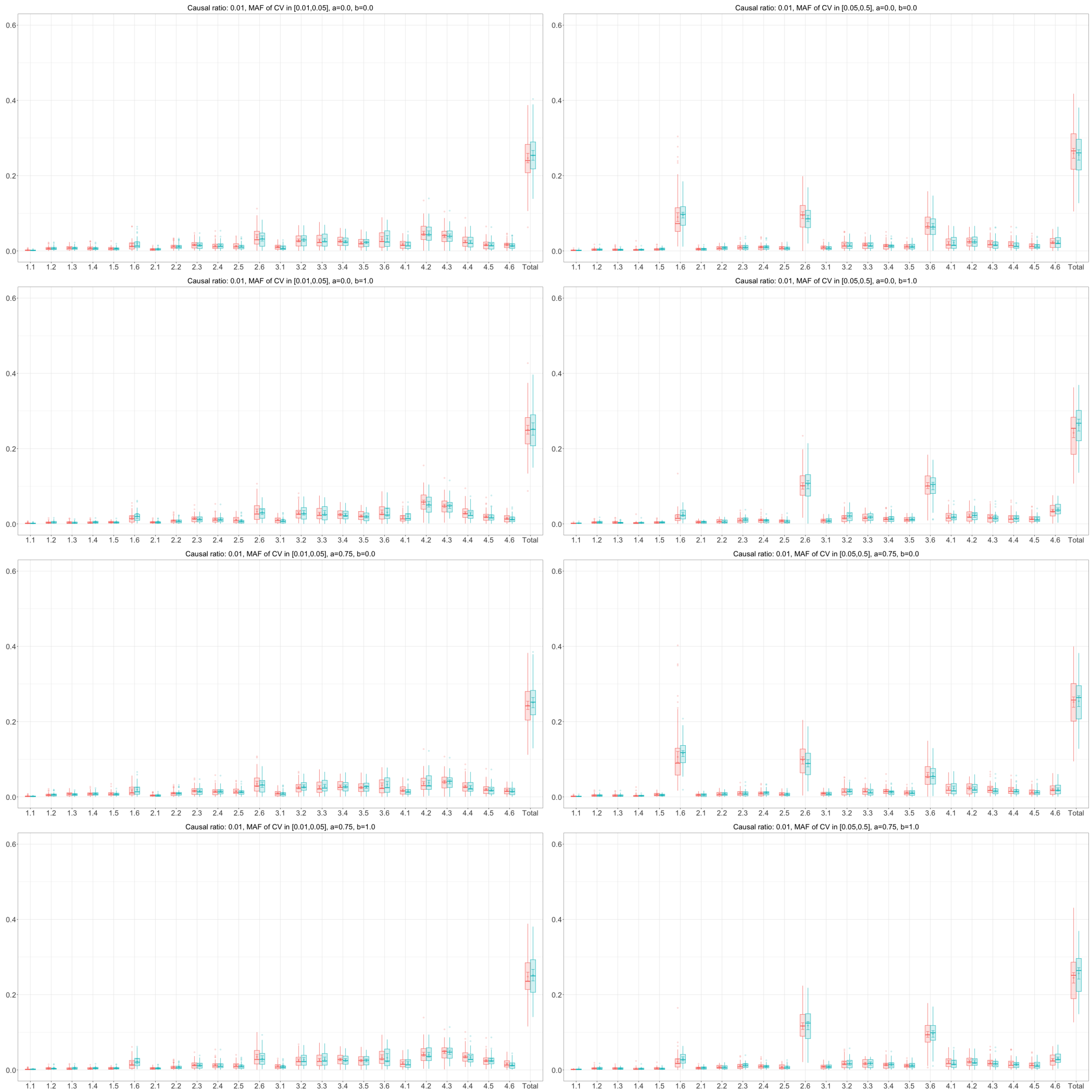

Figure S3: **Comparison of RHE-mc (red color) with GCTA-mc(blue color) in estimating partitioned heritability under 8 different genetic architectures on small-scale simulated data ($M = 590k$ array SNPs and $N = 10k$ individuals)**: We partition SNPs into 24 bins based on 6 MAF bins and 4 LDAK bins (Methods). True total SNP heritability is 0.25. Here x-axis represents the partitions ($i.j$ denotes the bin defined based on $i$-th ldak bin and $j$-th MAF bin. The lower bin number denotes the lower MAF (LDAK weights). For example, bin 1.6 contains SNPs which are in the first quartile of LDAK weights and MAF$> 0.05$ ). y-axis represents the heritability. Each boxplot shows the distribution of estimates from 100 simulations. Note that GCTA-mc did not run successfully on all 100 simulations.

| Genetic architecture | | Heritability (GCTA-mc) | | Heritability (RHE-mc) | |
|---|---|---|---|---|---|
| MAF of causal SNPs | MAF and LD coupling | Causal bin | Non-causal bin | Causal bin | Non-causal bin |
| $[0.01, 0.05]$ | $a = b = 0$ | 0.244±0.061 | 0.009± 0.051 | 0.242±0.064 | 0.004±0.052 |
| $[0.01, 0.05]$ | $a = 0, b = 1$ | 0.243±0.062 | 0.008±0.047 | 0.247±0.060 | 0.003±0.051 |
| $[0.01, 0.05]$ | $a = 0.75, b = 0$ | 0.241±0.061 | 0.009±0.050 | 0.240±0.062 | 0.002±0.051 |
| $[0.01, 0.05]$ | $a = 0.75, b = 1$ | 0.247±0.056 | 0.004± 0.048 | 0.244±0.06 | 0.003±0.051 |
| $[0.05, 0.5]$ | $a = b = 0$ | 0.251±0.048 | 0.012±0.003 | 0.251±0.052 | 0.007± 0.058 |
| $[0.05, 0.5]$ | $a = 0, b = 1$ | 0.248±0.052 | 0.014±0.054 | 0.240±0.049 | 0.001± 0.055 |
| $[0.05, 0.5]$ | $a = 0.75, b = 0$ | 0.255±0.047 | 0.000± 0.060 | 0.251±0.052 | 0.000± 0.060 |
| $[0.05, 0.5]$ | $a = 0.75, b = 1$ | 0.250±0.048 | 0.005±0.05 | 0.241±0.050 | 0.002±0.058 |

Table S3: **Heritability contribution of causal bin vs non-causal bins on small-scale simulated data ($M = 590k$ array SNPs and $N = 10k$ individuals)**: We ran both RHE-mc and GCTA-mc with 24 bins based on 6 MAF bins and 4 LDAK bins for 8 genetic architectures (Methods). In all simulations, the proportion of causal variants is 0.01 and true total heritability is 0.25. The causal SNPs are restricted to lie within a specific range of MAF, *i.e.*, within $[0.01, 0.05]$ for the first four rows and $[0.05, 0.5]$ for the last four. Non-causal bins refer to those bins where none of the SNPs is causal, *i.e.*, in each of the first four genetic architectures, these would correspond to bins with MAF $\notin [0.01, 0.05]$. Causal bins refer to all remaining bins. Standard errors are computed from 100 replicates.

#### S6.2 Heritability partitioning on large-scale data

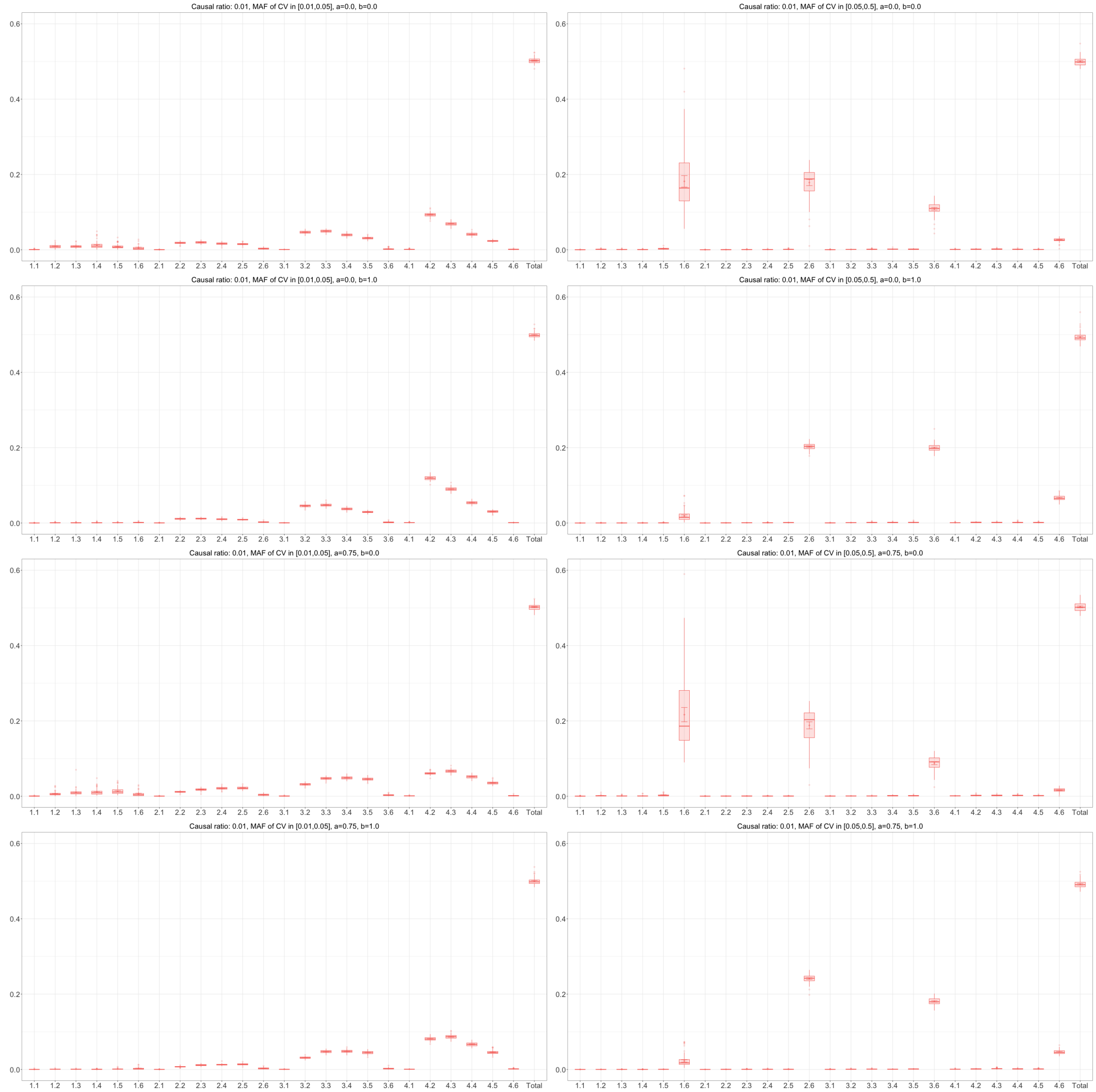

Figure S4: **Partitioned heritability estimates from RHE-mc on large-scale simulated data ($M = 590K$ array SNPs and $N = 337K$ individuals)**: We ran RHE-mc with 24 bins based on 6 MAF bins and 4 LDAK bins (Methods) over 8 different genetic architectures. Here x-axis represents the partitions ($i.j$ denotes the bin defined based on $i$-th ldak bin and $j$-th MAF bin) and y-axis represents the heritability. Each boxplot shows the distribution of estimates from 100 simulations.

| Genetic architecture | | | | Heritability | |
|---|---|---|---|---|---|
| Percentage of causal SNPs | MAF of causal SNPs | MAF and LD coupling | True total $h^2$ | Causal bin | Non-causal bin |
| 0.01 | [0.01,0.05] | a=b=0 | 0.5 | 0.501 ±0.006 | 0.000 ±0.004 |
| 0.01 | [0.01,0.05] | a=0,b=1 | 0.5 | 0.498 ±0.007 | 0.000 ±0.003 |
| 0.01 | [0.01,0.05] | a=0.75,b=0 | 0.5 | 0.500 ±0.008 | 0.002 ±0.004 |
| 0.01 | [0.01,0.05] | a=0.75,b=1 | 0.5 | 0.490 ±0.007 | 0.001 ±0.003 |
| 0.01 | [0.05,0.5] | a=b=0 | 0.5 | 0.501 ±0.036 | -0.001 ±0.030 |
| 0.01 | [0.05,0.5] | a=0,b=1 | 0.5 | 0.487 ±0.012 | 0.005 ±0.005 |
| 0.01 | [0.05,0.5] | a=0.75,b=0 | 0.5 | 0.508 ±0.026 | -0.005 ±0.023 |
| 0.01 | [0.05,0.5] | a=0.75,b=1 | 0.5 | 0.490 ±0.009 | 0.000 ±0.005 |

Table S4: **Heritability contribution of causal vs non-causal bins on large-scale simulated data ($M = 590K$ array SNPs and $N = 337K$ individuals)**: We ran RHE-mc with 24 bins based on 6 MAF bins and 4 LDAK bins (Methods). Standard errors are computed from 100 replicates.

#### S6.3 Accuracy of genome-wide SNP heritability

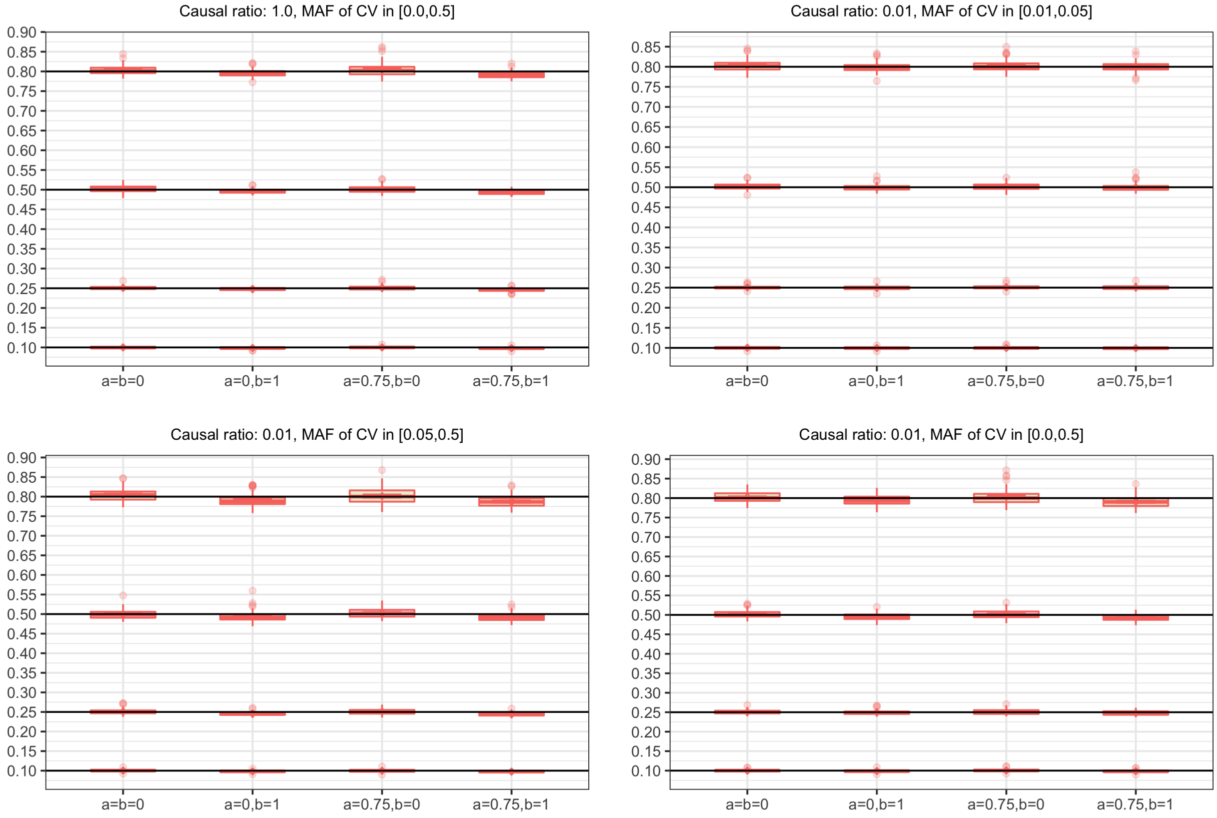

Figure S5: Accuracy of genome-wide SNP heritability estimated by RHE-mc across 64 distinct MAF- and LD-dependent architectures in genome-wide simulations ($N = 337,205$ unrelated individuals, $M = 593,300$ array SNPs). For simulating the phenotypes, we chose true heritability from $\{0.1, 0.25, 0.5, 0.8\}$, varied the ratio of causal variants (causal ratio $\in \{0.01, 1.0\}$), varied the MAF range of causal variants (MAF of CV), the coupling of MAF with effect size ($a = 0$ indicates no coupling of MAF and $a = 0.75$ indicates coupling of MAF), and the effect of local LD on effect size ($b = 0$ indicates no LDAK weights and $b = 1$ indicates LDAK weights) (see Methods). We ran RHE-mc using 24 bins formed by the combination of 6 bins based on MAF as well as 4 bins based on quartiles of the LDAK score of a SNP (see Methods). Each boxplot shows the distribution of RHE-mc estimates across 100 simulations.

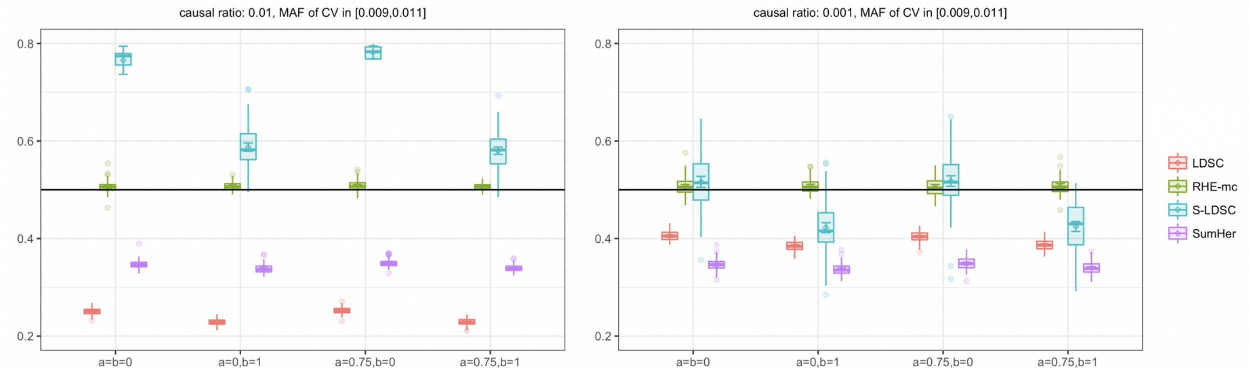

Figure S6: Comparison of estimates of genome-wide SNP heritability from RHE-mc with LDSC, S-LDSC , and SumHer when only rare variants are causal in large-scale simulations ($N = 337,205$ unrelated individuals, $M = 593,300$ array SNPs). We compared methods for heritability estimation under different genetic architectures when only rare variants are causal. We set true heritability to 0.5, the MAF range of causal variants (MAF of CV) to be between $[0.009, 0.011]$ and varied the coupling of MAF with effect size ($a = 0$ indicates no coupling of MAF and $a = 0.75$ indicates coupling of MAF), and the effect of local LD on effect size ($b = 0$ indicates no LDAK weights and $b = 1$ indicates LDAK weights) (see Methods). Here, we run RHE-mc using 24 bins formed by the combination of 6 bins based on MAF as well as 4 bins based on quartiles of the LDAK score of a SNP (see Methods). We run S-LDSC with 10 MAF bins (see Supplementary Table S5 ). To do a fair comparison, for every method, we computed LD scores and LDAK weights by using in-sample LD, and in all simulations we aim to estimate the SNP-heritability explained by the same set of M SNPs.

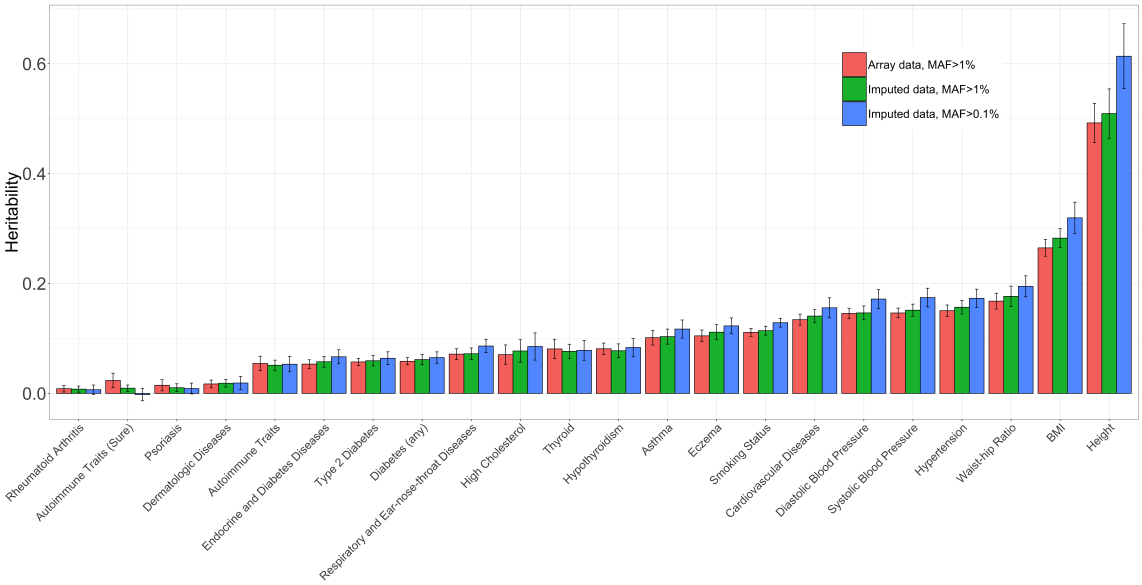

Figure S7: **Estimates of genome-wide SNP heritability from RHE-mc for 22 complex traits and diseases in the UK Biobank**: We restricted our analysis to $N = 291,273$ unrelated white British individuals. First, we applied RHE-mc to $M = 459,792$ array SNPs (MAF$>$ 1%) with 8 MAF/LD bins. Second, we applied RHE-mc to $M = 4,824,392$ imputed SNPs (MAF $>$ 1%) with 8 MAF/LD bins(Methods). Third, we applied RHE-mc to $M = 7,774,235$ imputed SNPs (MAF $>$ 0.1%) with 144 MAF/LD bins (Methods). Black bars mark $\pm 2$ standard errors.

| MAF bin | Range | Number of SNPs |
|---|---|---|
| 1 | [0, 0.0126) | 59330 |
| 2 | [0.0126, 0.020) | 59330 |
| 3 | [0.020, 0.029) | 59330 |
| 4 | [0.029, 0.0433) | 59330 |
| 5 | [0.043, 0.0658) | 59330 |
| 6 | [0.065, 0.106) | 59330 |
| 7 | [0.106, 0.170) | 59330 |
| 8 | [0.170, 0.260) | 59330 |
| 9 | [0.260, 0.373) | 59330 |
| 10 | [0.373, 0.5) | 59330 |

Table S5: MAF bins which are used in running S-LDSC over the large scale simulated data.

### S7 Parameter settings for summary statistics methods

For running LDSC we computed the LD score of each SNP within 2-Mb windows centered on the SNP. We ran LD score regression with an unconstrained intercept and with regression weights that account for correlations between association statistics at SNPs in LD and heteroscedasticity [2]. For preventing the LDSC software from dropping high-effect SNPs we used the following flags –not-M-5-50 and –chisq-max 99999.
In simulations, we ran S-LDSC with 10 binary MAF bins which are defined such that each bin contains 10% of the typed SNPs; this is done to reflect the 10 MAF bin annotations in the S-LDSC baseline-LD model [3] (see Table S5 for the details of MAF bins). In analysing the 22 real complex traits, we run S-LDSC with baseline-LD model[3].
To run SumHer, first we computed the default LDAK weights using in-sample LD [4]. After that we computed LD tagging using 1-Mb windows centered on each SNP and setting $\alpha = -0.25$ as recommended [5]. We used default values for the other parameter settings for running SumHer.

To do a direct comparison among LDSC, S-LDSC, and SumHer, we ran an in-sample LD version of each method meaning that we used same set of SNPs to compute LD scores and LDAK weights, perform the regression, and estimate SNP-heritability.

### S8 Continuous annotations

We assessed the accuracy of RHE-mc in estimating variance components with continuous annotation. We simulated a phenotype with true heritabillity 0.5 from $9K$ indviduals and $15k$ SNPs under the GCTA model. We ran RHE-mc with single component, no annotations, and standardized genotypes. We next ran RHE-mc with single component, non-standardized genotypes, where we added a continous annotation defined as $1/var(i)$ for SNP $i$ where $var(i)$ is the variance of SNP $i$ across individuals. We obtain concordant estimate of genome-wide SNP heritability $0.45 \pm 0.03$ in the first case and $0.46 \pm 0.03$ in the second case.

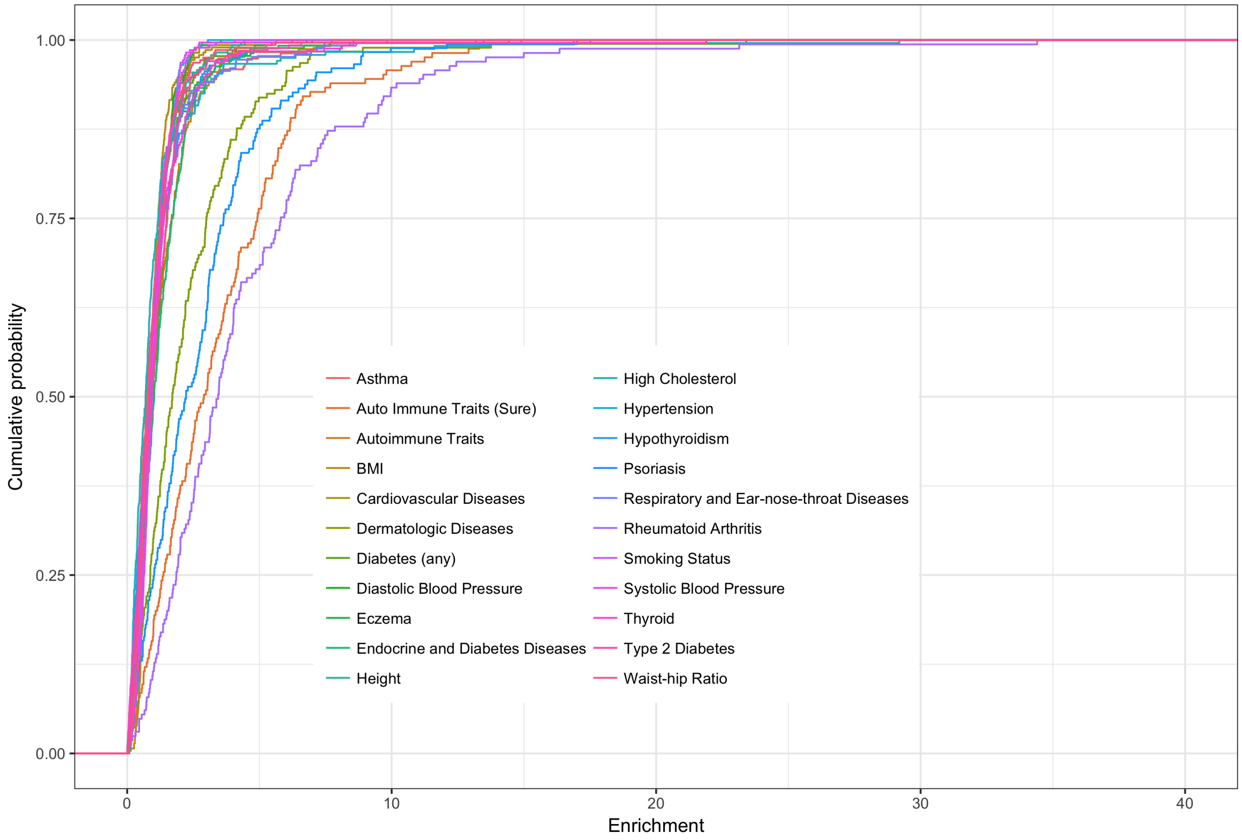

Figure S8: Partitioing of genome-wide SNP heritability from RHE-mc for 22 complex traits and diseases in the UK Biobank ($N = 291,273$ unrelated white British individuals, $M = 459,792$ common SNPs with respect to 300 bins defined based on 10M base pairs. Here we plot the empirical cumulative probability respectively of the enrichment.

### S9 Power as a function of annotation size

To quantify the power of RHE-mc as a function of the size of an annotation, we performed simulations using $N = 291,273$ unrelated white British individuals and $M = 459,792$ common SNPs. We defined 8 annotations (4 MAF bins and 2 LD bins) in which we fixed the heritability of a selected bin and varied the proportion of SNPs in the selected category. We then plotted probability of rejection; the results are displayed in Figure S10 . Furthermore, we simulated phenotypes in which we fixed the enrichment of a selected bins and varied the size of the selected bin, the results are displayed in Table S6.

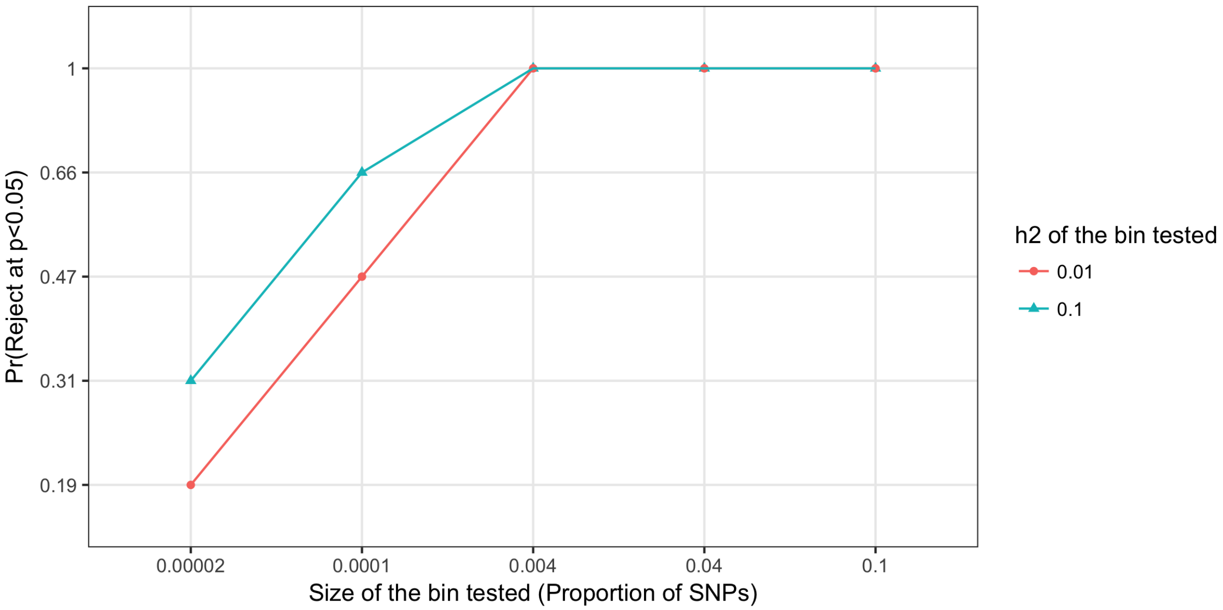

Figure S9: Power as a function of annotation size. Each point represents a rejection probability over 100 simulations. All simulations have $h^2_{total} = 0.7$, $N = 291,273$, $M = 459,792$, $p_{causal} = 0.05$.

| True enrichment | Proportion of SNPs | point estimate | SE | Pr(rejection at p< 0.05) |
|---|---|---|---|---|
| 2 | 0.4% | 2.06 | 0.4 | 100% |
| 1 | 0.4% | 1.02 | 0.14 | 100% |
| 0 | 0.4% | 0.0 | 0.02 | 0.5% |
| 2 | 0.01% | 2.18 | 1.07 | 30% |

Table S6: Power as function of annotation size. SE, point estimate, and probability of rejections are computed from 100 replicates. All simulations have $h^2_{total} = 0.7$, $N = 291,273$, $M = 459,792$, $p_{causal} = 0.05$.

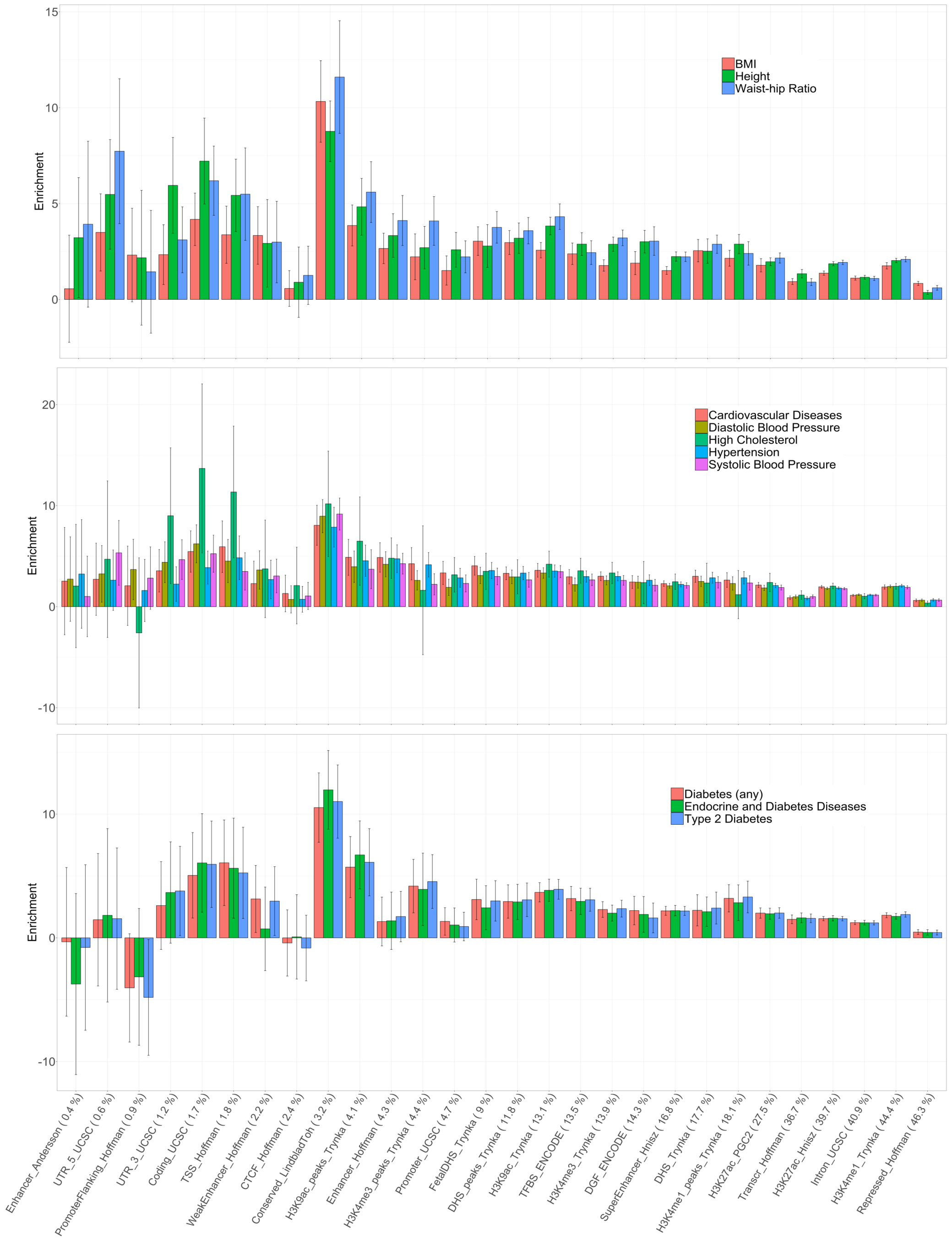
BMI
Height
Waist-hip Ratio
Enrichment
15
10
5
0
Cardiovascular Diseases
Diastolic Blood Pressure
High Cholesterol
Hypertension
Systolic Blood Pressure
Enrichment
20
10
0
-10
Diabetes (any)
Endocrine and Diabetes Diseases
Type 2 Diabetes
Enrichment
10
0
-10
Enhancer_Andersson ( 0.4 %)
UTR_5_UCSC ( 0.6 %)
PromoterFlanking_Hoffman ( 0.9 %)
UTR_3_UCSC ( 1.2 %)
Coding_UCSC ( 1.7 %)
TSS_Hoffman ( 1.8 %)
WeakEnhancer_Hoffman ( 2.2 %)
CTCF_Hoffman ( 2.4 %)
Conserved_LindbladToh ( 3.2 %)
H3K9ac_peaks_Trynka ( 4.1 %)
Enhancer_Hoffman ( 4.3 %)
H3K4me3_peaks_Trynka ( 4.4 %)
Promoter_UCSC ( 4.7 %)
FetalDHS_Trynka ( 9 %)
DHS_peaks_Trynka ( 11.8 %)
H3K9ac_Trynka ( 13.1 %)
TFBS_ENCODE ( 13.5 %)
H3K4me3_Trynka ( 13.9 %)
DGF_ENCODE ( 14.3 %)
SuperEnhancer_Hnisz ( 16.8 %)
DHS_Trynka ( 17.7 %)
H3K4me1_peaks_Trynka ( 18.1 %)
H3K27ac_PGC2 ( 27.5 %)
Transcr_Hoffman ( 36.7 %)
H3K27ac_Hnisz ( 39.7 %)
Intron_UCSC ( 40.9 %)
H3K4me1_Trynka ( 44.4 %)
Repressed_Hoffman ( 46.3 %)

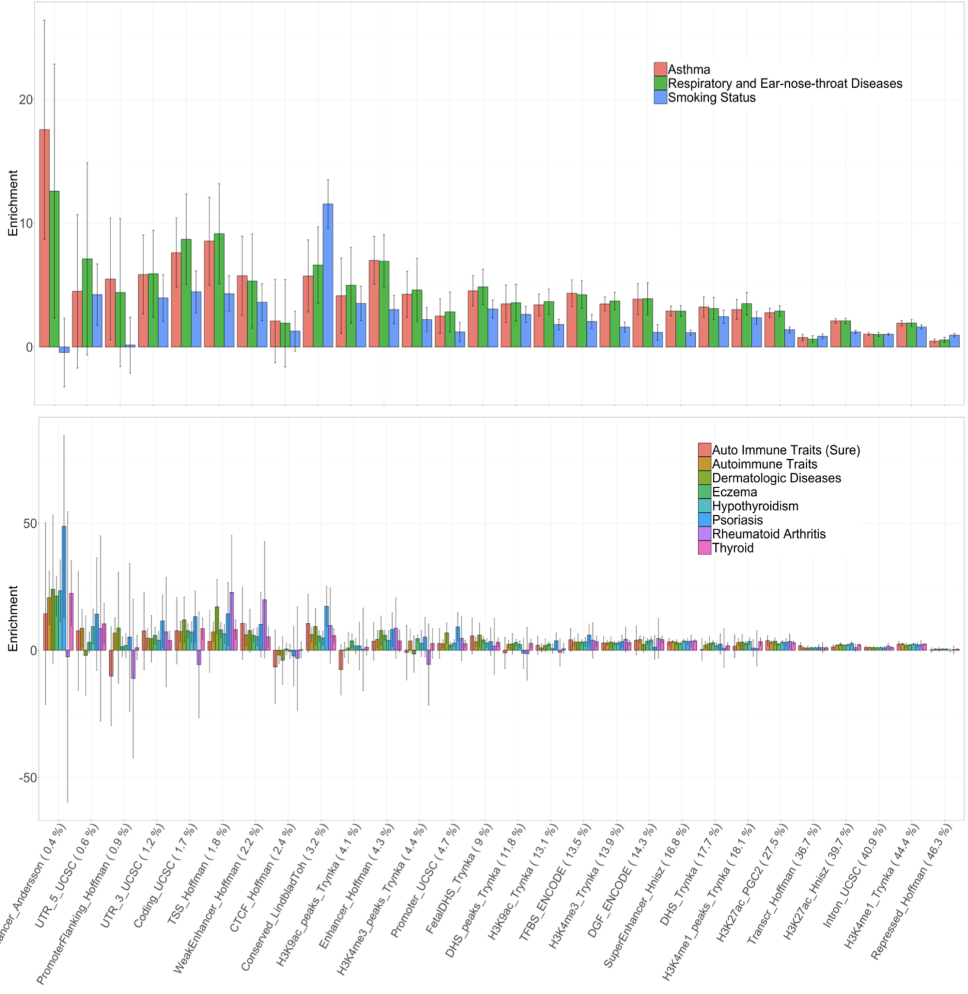

Figure S10: Enrichment of heritability across 28 functional annotations: We applied RHE-mc to $N = 291,273$ unrelated white British individuals and $M = 5,670,959$ imputed SNPs (MAF $> 0.1\%$ and present in 1000 Genomes Project). SNPs were partitioned based on 28 functional annotations that were defined in a previous study [6]. We grouped 22 traits in the UK Biobank into five categories (autoimmune, diabetes, respiratory, anthropometric, cardiovascular). Black bars mark $\pm 2$ standard errors. Annotations are ordered by the proportions of SNPs in that annotation (given in parentheses)

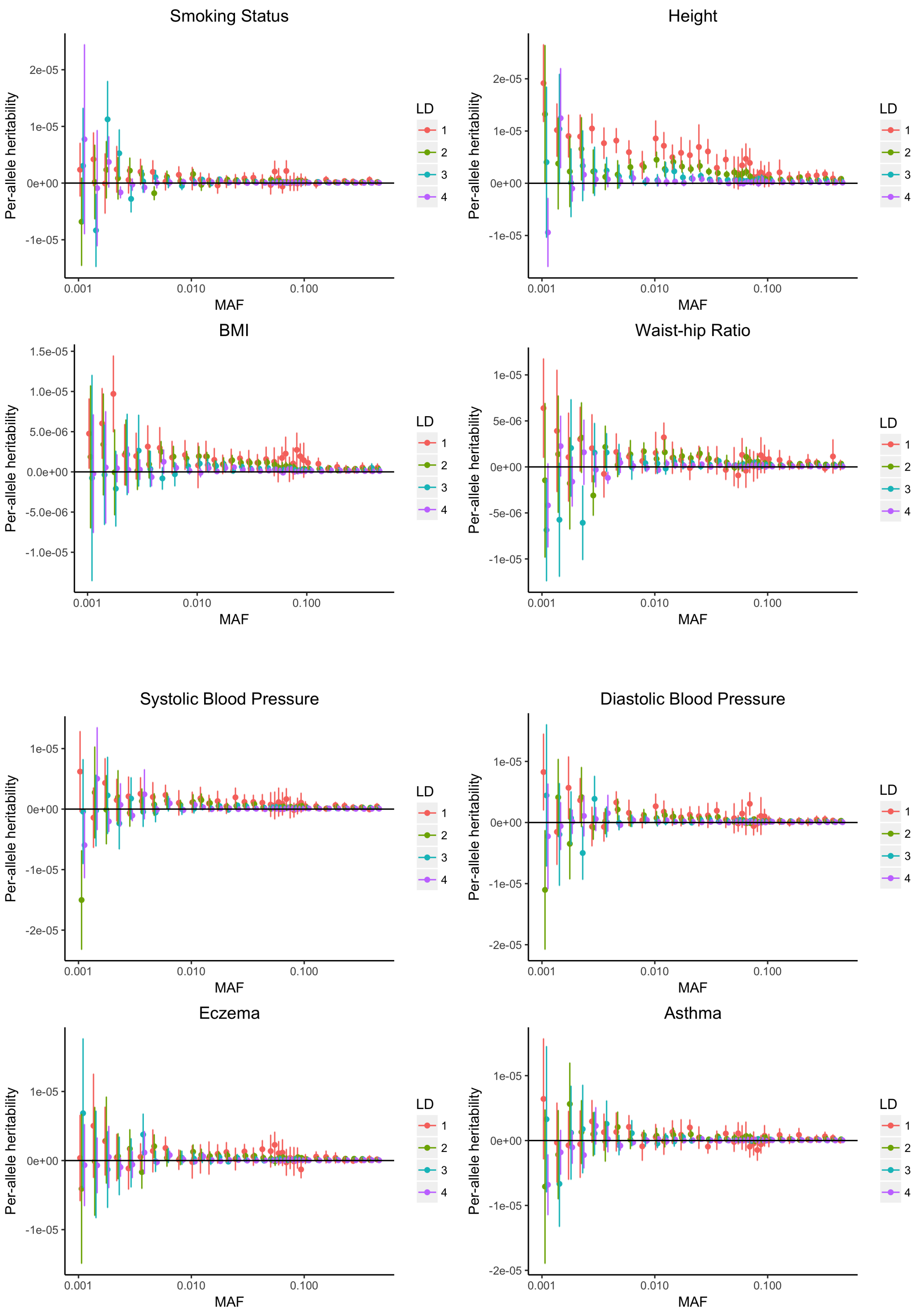
Smoking Status
Per-allele heritability
2e-05
1e-05
0e+00
-1e-05
0.001
0.010
0.100
MAF
LD
1
2
3
4
Height
Per-allele heritability
2e-05
1e-05
0e+00
-1e-05
0.001
0.010
0.100
MAF
LD
1
2
3
4
BMI
Per-allele heritability
1.5e-05
1.0e-05
5.0e-06
0.0e+00
-5.0e-06
-1.0e-05
0.001
0.010
0.100
MAF
LD
1
2
3
4
Waist-hip Ratio
Per-allele heritability
1e-05
5e-06
0e+00
-5e-06
-1e-05
0.001
0.010
0.100
MAF
LD
1
2
3
4
Systolic Blood Pressure
Per-allele heritability
1e-05
0e+00
-1e-05
-2e-05
0.001
0.010
0.100
MAF
LD
1
2
3
4
Diastolic Blood Pressure
Per-allele heritability
1e-05
0e+00
-1e-05
-2e-05
0.001
0.010
0.100
MAF
LD
1
2
3
4
Eczema
Per-allele heritability
1e-05
0e+00
-1e-05
0.001
0.010
0.100
MAF
LD
1
2
3
4
Asthma
Per-allele heritability
1e-05
0e+00
-1e-05
-2e-05
0.001
0.010
0.100
MAF
LD
1
2
3
4

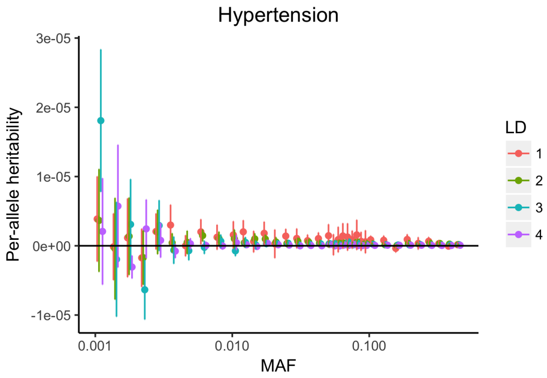
Hypertension
Per-allele heritability
3e-05
2e-05
1e-05
0e+00
-1e-05
0.001
0.010
0.100
MAF
LD
1
2
3
4

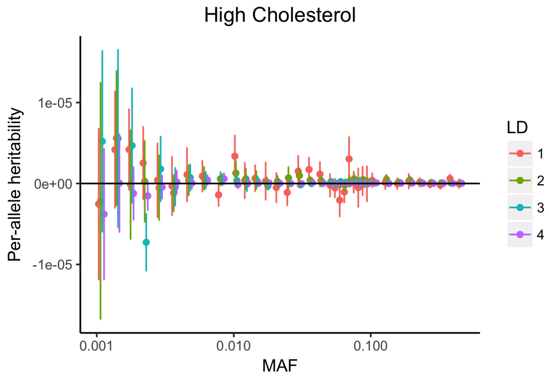
High Cholesterol
Per-allele heritability
1e-05
0e+00
-1e-05
0.001
0.010
0.100
MAF
LD
1
2
3
4

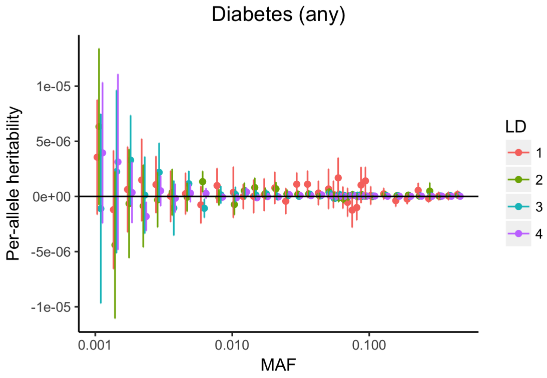
Diabetes (any)
Per-allele heritability
1e-05
5e-06
0e+00
-5e-06
-1e-05
0.001
0.010
0.100
MAF
LD
1
2
3
4

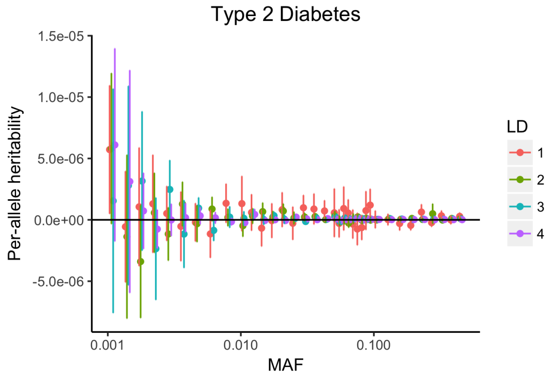
Type 2 Diabetes
Per-allele heritability
1.5e-05
1.0e-05
5.0e-06
0.0e+00
-5.0e-06
0.001
0.010
0.100
MAF
LD
1
2
3
4

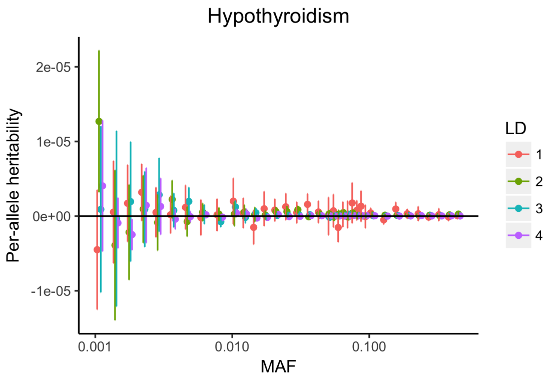
Hypothyroidism
Per-allele heritability
2e-05
1e-05
0e+00
-1e-05
0.001
0.010
0.100
MAF
LD
1
2
3
4

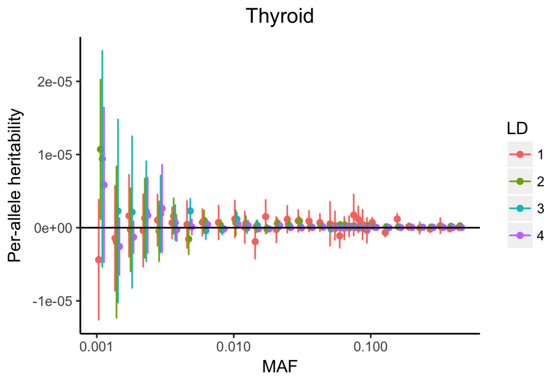
Thyroid
Per-allele heritability
2e-05
1e-05
0e+00
-1e-05
0.001
0.010
0.100
MAF
LD
1
2
3
4

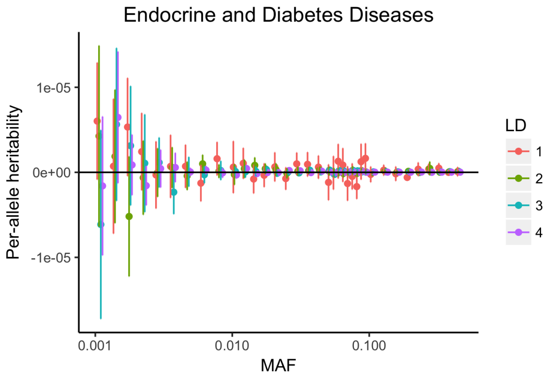
Endocrine and Diabetes Diseases
Per-allele heritability
1e-05
0e+00
-1e-05
0.001
0.010
0.100
MAF
LD
1
2
3
4

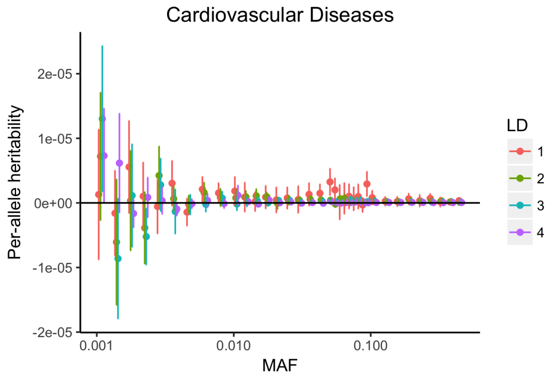
Cardiovascular Diseases
Per-allele heritability
2e-05
1e-05
0e+00
-1e-05
-2e-05
0.001
0.010
0.100
MAF
LD
1
2
3
4

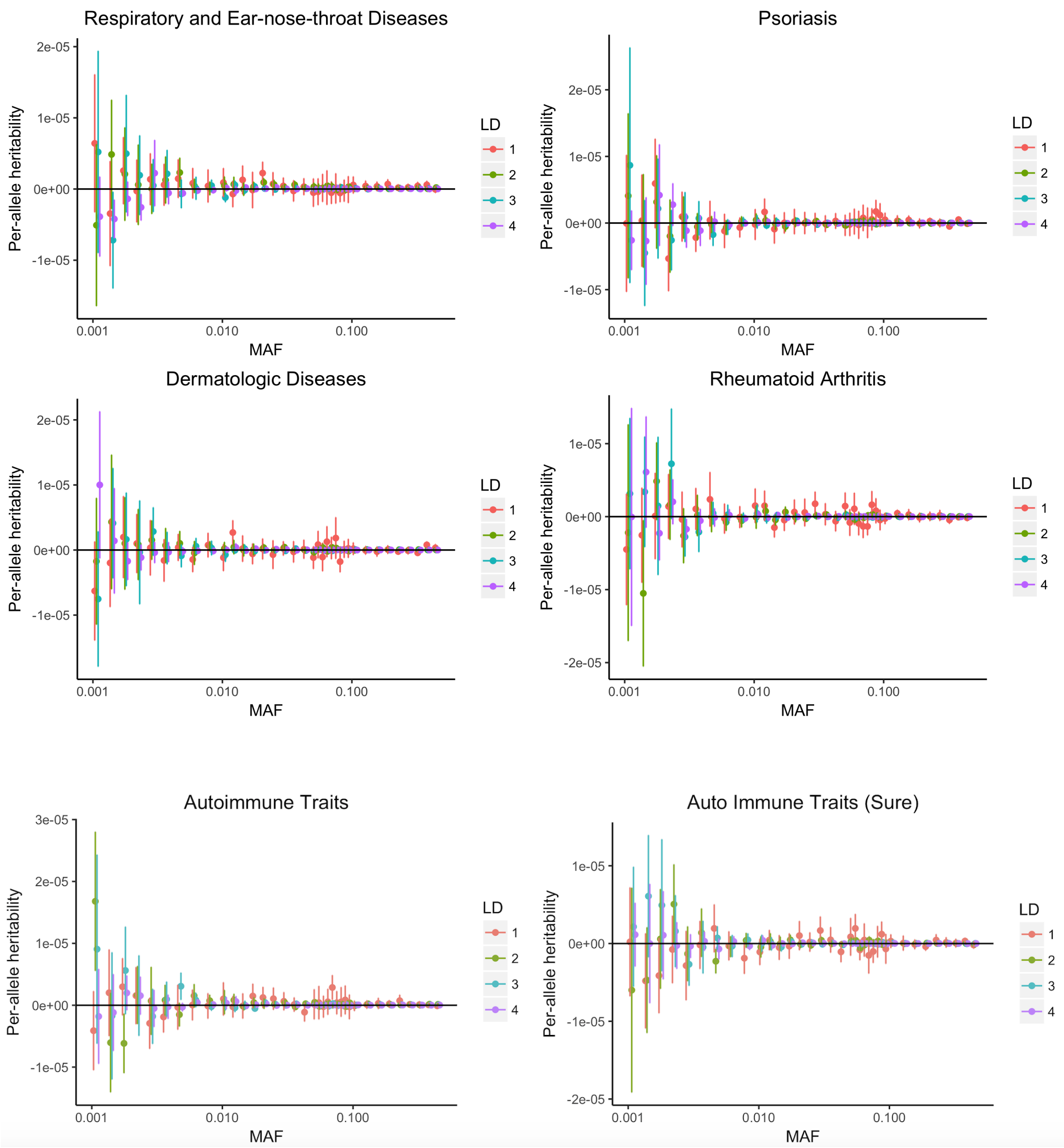

Figure S11: Per-allele heritability of 22 triats as a function of MAF: We applied RHE-mc to $N = 291,273$ unrelated white British individuals and $M = 7,774,235$ imputed SNPs. SNPs were partitioned into 144 bins based on LD score (4 bins based on quartiles of the LD score with $i$ denoting the $i^{th}$ quartile) and MAF (36 MAF bins) (see Methods). Per allele heritability for bin $k$ is defined as $\frac{h_k^2}{M_k * 2f_k * (1-f_k)}$ where $h_k^2$ is the heritability attributed to bin $k$, $M_k$ is the number of SNPs in bin $k$, and $f_k$ is the average MAF in bin $k$. Bars mark $\pm 2$ standard errors.

#### S10 Array UK Biobank data

| Trait | Heritability | | |
|---|---|---|---|
| | Chromosome | MAF/LD | 10Mb |
| Autoimmune Traits | $0.064 \pm 0.005$ | $0.054 \pm 0.006$ | $0.070 \pm 0.004$ |
| Auto Immune Traits (Sure) | $0.011 \pm 0.002$ | $0.023 \pm 0.006$ | $0.029 \pm 0.001$ |
| Dermatologic Diseases | $0.020 \pm 0.003$ | $0.0172 \pm 0.003$ | $0.021 \pm 0.001$ |
| Psoriasis | $0.017 \pm 0.002$ | $0.014 \pm 0.005$ | $0.022 \pm 0.003$ |
| Rheumatoid Arthritis | $0.008 \pm 0.002$ | $0.008 \pm 0.002$ | $0.010 \pm 0.003$ |
| Eczema | $0.124 \pm 0.007$ | $0.104 \pm 0.005$ | $0.13 \pm 0.006$ |
| Hypothyroidism | $0.097 \pm 0.008$ | $0.081 \pm 0.005$ | $0.11 \pm 0.007$ |
| Thyroid | $0.095 \pm 0.009$ | $0.081 \pm 0.008$ | $0.109 \pm 0.008$ |
| Diastolic Blood Pressure | $0.170 \pm 0.005$ | $0.145 \pm 0.004$ | $0.173 \pm 0.003$ |
| Systolic Blood Pressure | $0.172 \pm 0.006$ | $0.146 \pm 0.004$ | $0.171 \pm 0.004$ |
| Cardiovascular Diseases | $0.165 \pm 0.006$ | $0.134 \pm 0.005$ | $0.17 \pm 0.004$ |
| Hypertension | $0.179 \pm 0.006$ | $0.150 \pm 0.005$ | $0.183 \pm 0.006$ |
| High Cholesterol | $0.099 \pm 0.015$ | $0.070 \pm 0.008$ | $0.102 \pm 0.003$ |
| Diabetes (any) | $0.069 \pm 0.004$ | $0.058 \pm 0.003$ | $0.072 \pm 0.003$ |
| Endocrine and Diabetes Diseases | $0.064 \pm 0.004$ | $0.053 \pm 0.003$ | $0.065 \pm 0.003$ |
| Type 2 Diabetes | $0.068 \pm 0.004$ | $0.057 \pm 0.003$ | $0.069 \pm 0.005$ |
| BMI | $0.330 \pm 0.014$ | $0.264 \pm 0.007$ | $0.328 \pm 0.013$ |
| Height | $0.583 \pm 0.026$ | $0.492 \pm 0.017$ | $0.59 \pm 0.021$ |
| Waist-hip Ratio | $0.196 \pm 0.009$ | $0.167 \pm 0.007$ | $0.2 \pm 0.005$ |
| Asthma | $0.122 \pm 0.009$ | $0.101 \pm 0.006$ | $0.127 \pm 0.007$ |
| Smoking Status | $0.130 \pm 0.004$ | $0.111 \pm 0.003$ | $0.132 \pm 0.002$ |
| Respiratory and Ear-nose-throat Diseases | $0.086 \pm 0.007$ | $0.071 \pm 0.004$ | $0.091 \pm 0.004$ |

Table S7: Estimates of genome-wide SNP heritability from RHE-mc for 22 complex traits and diseases in the UK Biobank ($N = 291,273$ unrelated white British individuals, $M = 459,792$ common SNPs). We run RHE-mc with 8 bins defined based on two MAF bins (MAF$\leq$ 0.05, MAF$>$ 0.05) and quartiles of the LD-scores. Furthermore, we run RHE-mc with 22 bins defined based on chromosome number. On average, partitioning based on chromosome numbers leads 21% higher estimates of genome-wide SNP heritability for 22 traits than partitioning based on MAF and LD. For instance, it leads 18% and 13% higher estimates of heritability for height and BMI respectively. We also partitioned SNP based on 10 Mb genomic regions (300 variance components).
